# A phospho-switch in the N-terminus of NRT2.1 affects nitrate uptake by controlling the interaction of NRT2.1 with NAR2.1

**DOI:** 10.1101/2020.01.08.898254

**Authors:** Zhi Li, Xu Na Wu, Aurore Jaquot, Larence Lejay, Waltraud X Schulze

## Abstract

NRT2.1 can be phosphorylated at five different sites within N- and C-terminus. Here, we provide a systematic functional characterization of phosphorylation at S21 and S28 within the N-terminus of NRT2.1. We used existing phosphoproteomic data sets of nitrate starvation and nitrate resupply to construct a site-specific correlation network identifying kinase candidates to phosphorylate NRT2.1. By this approach, we identified NITRATE UPTAKE REGULATORY KINASE 1 (AT5G49770) which itself was regulated by phosphorylation at S839 and S870 within its kinase domain. In the active state, when S839 was dephosphorylated and S870 was phosphorylated, NURK1 was found to interact with NRT2.1 at dephosphorylated S28. Upon that interaction, NURK1 can phosphorylate NRT2.1 at S21. Phosphorylation of NRT2.1 at S21 resulted in low interaction of NRT2.1 with its activator protein NAR2.1. By contrast, phosphorylation of NRT2.1 at S28 by a yet unknown kinase enhanced the interaction with NAR2.1, but inhibited the interaction with NURK1. We propose that serines S21 and S28 are involved in a phospho-switch mechanism and by which the interaction of NRT2.1 with its activator NAR2.1, and thus NRT2.1 activity, is modulated. NURK1 here was identified as the kinase affecting this phospho-switch through phosphorylation of NRT2.1 at S21 leading to inactivation of NRT2.1.

## Introduction

Nitrate is the main form of nitrogen uptake for most plant growing in temperate regions. Nitrate is taken up from the soil by members of the NRT2 and NPF protein families (Leran et al., 2014). The uptake of nitrate through dedicated transporters has been under study for a long time (Krapp et al., 2014), and has identified several nitrate transporters in roots as well as in green tissue (Kiba et al., 2012). These nitrate transporters can be classified into two major groups based on their uptake modes as a low affinity transport system (LATS) or a high affinity transport system (HATS) (Wang et al., 2012). The LATS works with affinities for nitrate in the millimolar range, while HATS works with affinities for nitrate in micromolar range.

NRT2.1 (AT1G08090) was found to be the major high affinity nitrate transporter in Arabidopsis roots as shown by the loss of 75% of HATS activity in *nrt2.1* knock-out mutants (Cerezo et al., 2001). A strong correlation between root nitrate influx and *NRT2.1* gene expression was observed (Lejay et al., 1999; Girin et al., 2007), leading to the conclusion that transcriptional regulation of NRT2.1 is a key mechanism to control nitrate uptake under high affinity conditions. Consequently, the regulation of NRT2.1 in the past has mainly been studied at the gene transcriptional level. *NRT2.1* gene expression was shown to be induced upon nitrate supply (Lejay et al., 1999) and repressed by high nitrogen metabolism and high external nitrate concentrations (Lejay et al., 1999; Girin et al., 2007). In contrast, *NRT2.1* gene expression was up-regulated by light and sugars (Lejay et al., 1999; Lejay et al., 2003) and in general is well known to be affected by the C/N status of the plant (Girin et al., 2007; Krouk et al., 2010). Reciprocal regulation of NRT2.1 and ammonium transporters upon nitrate and ammonium influx was described (Gansel et al., 2001; Camanes et al., 2012).

In addition, NRT2.1 is well known to be regulated by protein-protein interaction with a small single-transmembrane protein, NAR2.1 (Orsel et al., 2007; Laugier et al., 2012). Both proteins need to be present in the plasma membrane to result in full nitrate uptake activity (Orsel et al., 2006), and particularly, *nar2.1* mutants seem to be defective in the trafficking of NRT2.1 to the plasma membrane (Wirth et al., 2007). NAR2.1 was proposed to form a hetero-oligomer with NRT2.1 (Yong et al., 2010), but the precise mechanism by which this complex formation is regulated remains unclear. Furthermore, the abundance of NRT2.1 protein within the membrane remains rather constant despite measureable changes in HATS activity upon stimulation by high nitrate, sugars, or light (Wirth et al., 2007; Laugier et al., 2012), suggesting further tight control of NRT2.1 activity by posttranslational modifications. Posttranslational control of nitrate transport activity is well known for dual-affinity transporter NRT1.1 (Liu et al., 1999; Guo et al., 2001): Phosphorylation at threonine 101 acts like a switch: When T101 is phosphorylated, NRT1.1 (AT1G12110) acts as a high affinity transporter. When T101 is dephosphorylated, NRT1.1 acts as a low affinity transporter (Liu and Tsay, 2003). This affinity switch by phosphorylation has recently been related to structural changes in characteristic domains of NRT1.1 (Parker and Newstead, 2014), and CIPK23 (AT1G30270) has been identified as a kinase phosphorylating NRT1.1 at T101 (Ho et al., 2009). Posttranslational control is also encountered in the NH_4_ ^+^transporter AMT1.1 that is phosphorylated at a C-terminal threonine residue in an NH_4_ ^+^concentration and time-dependent manner. This phosphorylation leads to allosteric inactivation of the transporter, down regulating NH_4_ ^+^transport to prevent NH_4_ ^+^toxicity (Loque et al., 2007; Lanquar et al., 2009), and is even catalyzed by the same kinase, CIPK23 (Straub et al., 2017).

With accumulation of global proteomics data sets acquired under nitrate stimulation (Engelsberger and Schulze, 2012; Wang et al., 2013) or nitrate deprivation (Menz et al., 2016), several phosphorylation sites for NRT2.1 were identified. For example, resupply of 3 mM nitrate to nitrate starves seedlings lead to a rapid dephosphorylation of NRT2.1 at S28 (Engelsberger and Schulze, 2012). However, the functional consequence of NRT2.1 phosphorylation is yet unclear, and the mechanism by which the phosphorylation level of NRT2.1 is modulated, i.e. the respective kinases and phosphatases remain uncharacterized.

In this work we performed a systematic study of the NRT2.1 posttranslational modification status under different conditions of nitrate availability and aimed to identify kinases that might excel posttranslational control of the major nitrate uptake transporter NRT2.1.

## Materials and Methods

### Experimental design and statistical rationale

Two phosphoproteomics data set were used to study NRT2.1 phosphorylation and reconstruction of kinase-substrate networks as described in Results section. In all experiments, only root tissue was analyzed. Firstly, we used a nitrate deprivation data set (Supplementary Figure S1A) in which plants were grown in hydroponic culture at 3 mM nitrate and then subjected to nitrate deprivation by transferring the plants to nitrate-free nutrient solution. Harvests were carried before transfer to nitrate-free solution (0 minutes), and after 15 minutes and 3 hours of nitrate deprivation. This data set was published previously (Menz et al., 2016). Secondly, we conducted a nitrate-resupply experiment (Supplementary Figure S1B), in which plants were grown at 3 mM nitrate, then starve for nitrate for two days. After starvation, nitrate was resupplied at a concentration of 0.2 mM or 5 mM for 5 and 15 minutes. In all experiments, at least three biological replicates of root tissue were processed for (phospho)proteomic analysis and results are presented as averages with standard deviation. Statistical comparisons were carried out by pairwise t-tests (two sample comparisons) or ONEWAY ANOVA (multiple sample comparisons).

### Plant material

Most experiments were carried out with Arabidopsis thaliana wild type (col-0). Furthermore, the following homozygous knock-out mutants were used: *nrt2.1-2* (Cerezo et al., 2001) and *at5g49770* (SALK_068035).

### Constructs

All constructs were made using Gateway technology (Invitrogen) according to the manufacturer’s instructions. For ratiometric bimolecular Fluorescence Complementation (rBiFC) of Arabidopsis proteins, cDNAs of the following genes were cloned into rBiFC plasmids (Grefen and Blatt, 2012): AT5G49770 with phosphorylation site mutations at T792, S839, S870, S919 and NRT2.1 with phosphorylation site mutations at S21 and S28. Phosphorylation sites were mutated either to alanine (no phosphorylation) or aspartate (putative phosphorylation mimic). For purification of the cytoplasmic domain of AT5G49770 protein to be used in *in vitro* kinase assays, the cytoplasmic domains were cloned into *Escherichia coli* BL21(DE3) expression plasmid pETGST 1a and fused with His and GST tags (plasmid Plasmids His-GST-AT5G49770-CD). Cytoplasmic domains of two homologs (AT5G49760 and AT5G49780) were also used.

### Protein expression and purification

Plasmids His-GST-AT5G49770-CD with and without respective phosphorylation site mutations, as well as His-GST-AT5G49760-CD and His-GST-AT5G49780-CD were transformed into *Escherichia coli* BL21 (DE3). After 5 h induction by IPTG (isopropyl ß-D-thiogalactopyranoside), cells were harvested and lysed using BugBuster Protein Extraction Reagent (Novagen, Nottingham, UK), soluble fractions were used over gravity flow Ni^2+^-NTA sepharose columns (1 ml, IBA GmbH, Goettingen, Germany).

### In vitro kinase activity assay

In vitro kinase assays were performed as described in ADP-Glo™ Kinase assay kit (Promega, Germany) with minor modifications. 1 nmol kinase (cytoplasmic domain) was incubated with 5ug substrate in kinase reaction buffer (40mM Tris-HCl, pH 7.5, 1mM MgCl2, 0,01%BSA, 50 uM NaF, 1 uM Na3VO3, 1mM CaCl2, 100 uM ATP, 1 mM DTT). NRT2.1 peptides obtained from the QconCAT artificial protein were used as substrates. After incubation at room temperature for 1 hour, reaction was be terminated by heating at 90°C for 10 minutes. Then phosphopeptides were collected over Titanium dioxide as described above and analyzed by LC-MS/MS.

A luciferase-based kinase activity assay was performed according to the manufacturer’s instructions of ADP-Glo™ Kinase assay kit (Promega, Germany). 1nmol kinase (cytoplasmic domain) was incubated with generic kinase substrate myelin basic protein in kinase reaction buffer. After incubation for one hour, 30 µl ADP-GLO Reagents (Promega, Germany) was added and incubated for 40 minutes. Then Kinase Detection Reagents were added and incubated for another hour. Luminescence as a measure of ATP conversion from ADP was recorded with a luminometer (Tecan M200 Pro).

### Ratiometric bimolecular fluorescence assay

Positive colonies of *Agrobacterium tumefaciens* harboring the relevant constructs described above were propagated in LB medium at 28°C for 2 days, and then diluted as 1:500 into new medium for overnight culture. *Agrobacterium* pellets were collected by centrifugation at 1500 g for 15 min and resuspended in resuspension buffer (10 mM MES, 10 mM MgCl_2_, 0.15mM acetosyringone, pH 5.8), the suspension was diluted to OD600 of 0.5. The suspensions were injected into 5 to 6 weeks old *Nicotiana benthamiana* leaves with 1 ml syringe for 2 days before observation. Fluorescence was observed using a Zeiss LSM700 confocal microscope (20X 0.75-NA objectives). In all cases, excitation intensities, filter settings, photomultiplier gains and other parameters were standardized. The YFP and RFP fluorochromes were excited with 488 nm and 561 nm, respectively. Emitted light was collected at a range of 500-560nm for YFP and 575-625 nm for RFP. All images throughout all experiments were collected using exactly the same settings. The collected images were processed and both YFP and RFP intensity was measured using the FIJI software (Schindelin et al., 2012) and the YFP/RFP ratio was calculated. 10-50 different cells from leaves of the 2 plants were analyzed. Statistical significance was determined using Welch Two Sample t-test. To calibrate YFP/RFP ratios, the known interaction of NRT2.1 with NAR2.1 was used as reference (Laugier et al., 2012).

### Nitrate influx assay

Root NO_3_ influx was assayed as described (Laugier et al., 2012). Briefly, the plants were sequentially transferred to 0.1 mM CaSO_4_ for 1 min, to a complete nutrient solution, pH 5.8, containing 0.2mM ^15^NO_3_ (99 atom % excess ^15^N) for 5 min, and finally to 0.1mM CaSO_4_ for 1 min. Roots were then separated from shoots, and the roots dried at 70 °C for 48 h. After determination of their dry weight, the samples were analyzed for total nitrogen and atom % ^15^N using a continuous flow isotope ratio mass spectrometer coupled with a C/N elemental analyzer (model ANCA-MS; PDZ Europa, Crewe, UK). Each influx value is the mean of 6 to 12 replicates.

### Gene expression analysis

Root samples were frozen in liquid N_2_ in 2mL tubes containing one steel bead (2.5 mm diameter). Tissues were disrupted for 1 min at 28 s^-1^ in a mixer mill homogenizer (MM301, Retsch, Germany). Total RNA was extracted from tissues using TRizol reagent (Invitrogen, Carlsbad, CA, USA). Subsequently, 4 µg of RNAs were treated with DNase I (Amplification Grade, Sigma-Aldrich) following the manufacturer’s instructions. Reverse transcription was achieved with 4 µg of RNAs in the presence of Moloney Murine Leukemia Virus (M-MLV) reverse transcriptase (RNase H minus, Point Mutant, Promega) after annealing with an anchored oligo(dT)18 primer as described (Wirth et al., 2007). The quality of the cDNA was verified by PCR using specific primers spanning an intron in the gene APTR (At1g27450) forward 5’-CGCTTCTTCTCGACACTGAG-3’; reverse 5’-CAGGTAGCTTCTTGGGCTTC-3’.

Gene expression was determined by quantitative real-time PCR (LightCycler 480, Roche Diagnostics) using the SYBRR Premix Ex TaqTM (TaKaRa) according to the manufacturer’s instructions with 1 µl of cDNA in a total volume of 10 µl. The conditions of amplifications were performed as described previously (Wirth et al., 2007), except the first 10 minutes at 95°C which has become 30 seconds at 95°C. All the results presented were standardized using the housekeeping gene Clathrin (At4g24550). Gene-specific primer sequences were: NRT2.1 forward, 5’-AACAAGGGCTAACGTGGATG-3’; NRT2.1 reverse, 5’-CTGCTTCTCCTGCTCATTCC-3’; NAR2.1 forward, 5’-GGCCATGAAGTTGCCTATG -3’; NAR2.1 reverse, 5’-TCTTGGCCTTCCTCTTCTCA -3’; Clathrin forward, 5’-AGCATACACTGCGTGCAAAG-3’; Clathrin reverse, 5’-TCGCCTGTGTCACATATCTC-3’. At5g49770 forward 5’-CCAACCGTAACTTGAAAGGAAAGC-3’ At5g49770 reverse 5’-TCAGGATTGCCAGTCAAATCCAAG-3’.

### Microsomal protein enrichment

A total of 1 to 1.5 g of roots (fresh weight) was homogenized in 10 ml ice-cold extraction buffer (330mM mannitol, 100 mM KCl, 1 mM EDTA, 50 mM Tris-MES, fresh 5 mM DTT, and 1 mM phenylmethylsulfonylfluoride, pH 7.5) (Pertl et al., 2001) in the presence of 0.5% v/v proteinase inhibitor mixture (Sigma-Aldrich, Germany) and phosphatase inhibitors (25 mM NaF, 1 mM Na3VO4, 1 mM benzamidin, 3 µM proteinase inhibitor leupeptin). The homogenate was centrifuged for 15 minutes at 7500 × g at 4°C. The pellet was discarded, and the supernatant was centrifuged for 75 minutes at 48,000 × g at 4°C. Microsomal pellets were stored at −80°C until further processing.

### Protein digestion and phosphopeptide enrichment

The microsomal pellet was resuspended in 100 µl of membrane buffer (330 mM mannitol, 25 mM Tris-MES, 0.5 mM DTT) or UTU (6 M urea, 2 M thiourea, pH 8). Further tryptic digestion, desalting over C18 and enrichment of phosphopeptides over titanium dioxide beads was performed as described (Wu et al., 2017).

### LC-MS/MS analysis of peptides

Peptides mixtures were analyzed by nanoflow Easy-nLC (Thermo Scientific) and Orbitrap hybrid mass spectrometer (Q-exactive, Thermo Scientific). Peptides were eluted from a 75 µm x 50 cm analytical C_18_ column (PepMan, Thermo Scientific) on a linear gradient running from 4% to 64% acetonitrile over 135 min. Proteins were identified based on the information-dependent acquisition of fragmentation spectra of multiple charged peptides. Up to twelve data-dependent MS/MS spectra were acquired in the linear ion trap for each full-scan spectrum acquired at 70,000 full-width half-maximum (FWHM) resolution.

### Protein identification and label-free quantitation

MaxQuant version 1.5.3.8 (Cox and Mann, 2008) was used for raw file peak extraction and protein identification against the UniProt Arabidopsis database UP000006548 (39,389 entries). Protein quantification was performed in MaxQuant using the label free quantification (LFQ) algorithm (Cox et al., 2014). The following parameters were applied: trypsin as cleaving enzyme; minimum peptide length of seven amino acids; maximal two missed cleavages; carbamidomethylation of cysteine as a fixed modification; N-terminal protein acetylation, oxidation of methionine as variable modifications. For phosphopeptide identification also the phosphorylation of serine, threonine, and tyrosine was included as variable modifications. Peptide mass tolerance was set to 20 ppm and 0.5 Da was used as MS/MS tolerance. Further settings were: “label-free quantification” marked, multiplicity set to 1; “match between runs” marked with time window 2 min; peptide and protein false discovery rates (FDR) set to 0.01; common contaminants (trypsin, keratin, etc.) excluded. Phosphorylation sites were determined by the site-scanning algorithm search engine Andromeda (Cox et al., 2011). Phosphopeptide identifications were submitted to the PhosPhAt database (Durek et al., 2010). The raw MS data from this study was deposited at the ProteomeXchange Consortium (http://proteomecentral.proteomexchange.org) via the PRIDE partner repository with the identifier PXD015390 for the nitrate starvation data set, and PXD014146 for the nitrate resupply data set.

### Statistical analyses and data visualization

Phosphosites data (Phospho(STY)Sites.txt) were analyzed by Perseus software (Tyanova et al., 2016). Briefly, phosphosites quantified in at least 50% in all of samples were analyzed by ANOVA. Other statistical analyses were carried out with Sigma Plot (version 11.0) and Excel (Microsoft, 2013). Over-representation analysis was done via Fisher’s exact test, p values were adjusted using Bonferroni correction.

### Public resources

Functional classification of proteins was done based on MAPMAN (Thimm et al., 2004). Information about subcellular location was derived from SUBA3 (Tanz et al., 2013). Detailed protein function was manually updated with the support of TAIR (Huala et al., 2001). Published protein-protein interactions were retrieved from STRING (Franceschini et al., 2013).

## Results

This work is based on two large-scale phosphoproteomics data sets of wild type roots describing phosphorylation events induced by nitrate deprivation (Menz et al., 2016) (Supplementary Figure S1A), and nitrate resupply (supplementary figure S1B). The mass spectrometry data of these two experimental sets were jointly processed and quantified. Thus, in the combined data set, 4746 phosphopeptides were identified (Supplementary Table S1), and at least one quantitative values was obtained for 4670 phosphopeptides. Nitrate deprivation and nitrate resupply affected the regulation of diverse processes at the plasma membrane, particularly the activity of plasma membrane H^+^-ATPases (bin 34.1), and aquaporins (bin 34.19) which were down-regulated by nitrate-starvation and up-regulated by nitrate resupply. Protein synthesis (bin 29.2.1) and various signaling pathways through receptor kinases (bin 30.2) and calcium (bin 30.3) were also affected by both nitrate treatments. Glycolysis (bin 4.1) was only affected by nitrate deprivation (Supplementary Figure S1C), while nitrate assimilation (bin 12.1.1) was only affected (up-regulated) by nitrate resupply (Supplementary Figure S1C).

NRT2.1 was found to be phosphorylated at different sites in the N- and C-terminus, namely at S11 ((ac)GDSTGEPGSS(ph)MHGVTGR), S28 (EQSFAFSVQS(ph)PIVHTDK), S501 (NMHQGS(ph)LR), and T521 (SAAT(ph)PPENTPNNV) with distinct phosphorylation levels in nitrate starvation or nitrate resupply. Phosphorylation at S28 was identified already in previous starvation-resupply experiments (Engelsberger and Schulze, 2012). The phosphopeptides corresponding to phosphorylation at S11, S28, S501 and T521 sites were also identified in independent nitrate nutrition experiments (https://www.biorxiv.org/content/10.1101/583542v1). Phosphorylation at S21 with peptide EQS(ph)FAFSVQSPIVHTDK (Supplementary Figure S2, Supplementary Table S1) has not been explicitly described previously and was here firstly identified by careful re-analysis of nitrate deprivation data sets (Menz et al., 2016). The peptide and was also found in separately conducted experiments with Arabidopsis roots using resupply of 0.2 mM and 5 mM nitrate to stimulate high and low affinity uptake systems, respectively.

### NRT2.1 activity is regulated by N-terminal phosphorylation

To understand the role of different N-terminal phosphorylation sites for the activity of NRT2.1, site directed mutants of NRT2.1 were created, in which phosphorylation sites were mutated to a putative phosphomimicking aspartate (D), or phosphodead alanine (A). These phosphorylation site mutated NRT2.1 versions were expressed under the *NRT2.1* promoter in the *nrt2.1* knock-out background, and nitrate influx was measured for nitrate starved plants, as well as for plants under nitrate induction conditions with 1mM nitrate resupply for 1 and 4 hours. In wild type, higher nitrate influx was measured at 4 hours of induction by 1mM nitrate compared to the nitrate influx rates of roots at nitrate starvation. In the *nrt2.1-2* knock-out mutant, no induction of nitrate uptake was observed by 1mM nitrate treatment, resulting in significantly lower nitrate uptake compared to wild type after 4 h of induction with 1mM nitrate (Figure 1). For phosphorylation site mutations at NRT2.1 S11 we observed wild-type like influx of ^15^NO_3_ under all conditions in the phosphodead S11A mutant, while nitrate influx was significantly reduced in phosphomimicking S11D mutants (Figure 1A). This suggests that phosphorylation of NRT2.1 at S11 inhibits nitrate influx. For phosphorylation site mutations at S28 we observed and significantly reduced nitrate influx for NRT2.1S28A mutants, and wild-type like nitrate influx for phosphomimicking mutation NRT2.1S28D (Figure 1B). This suggests that phosphorylation of NRT2.1 at S28 activates nitrate influx.

**Figure 1:**
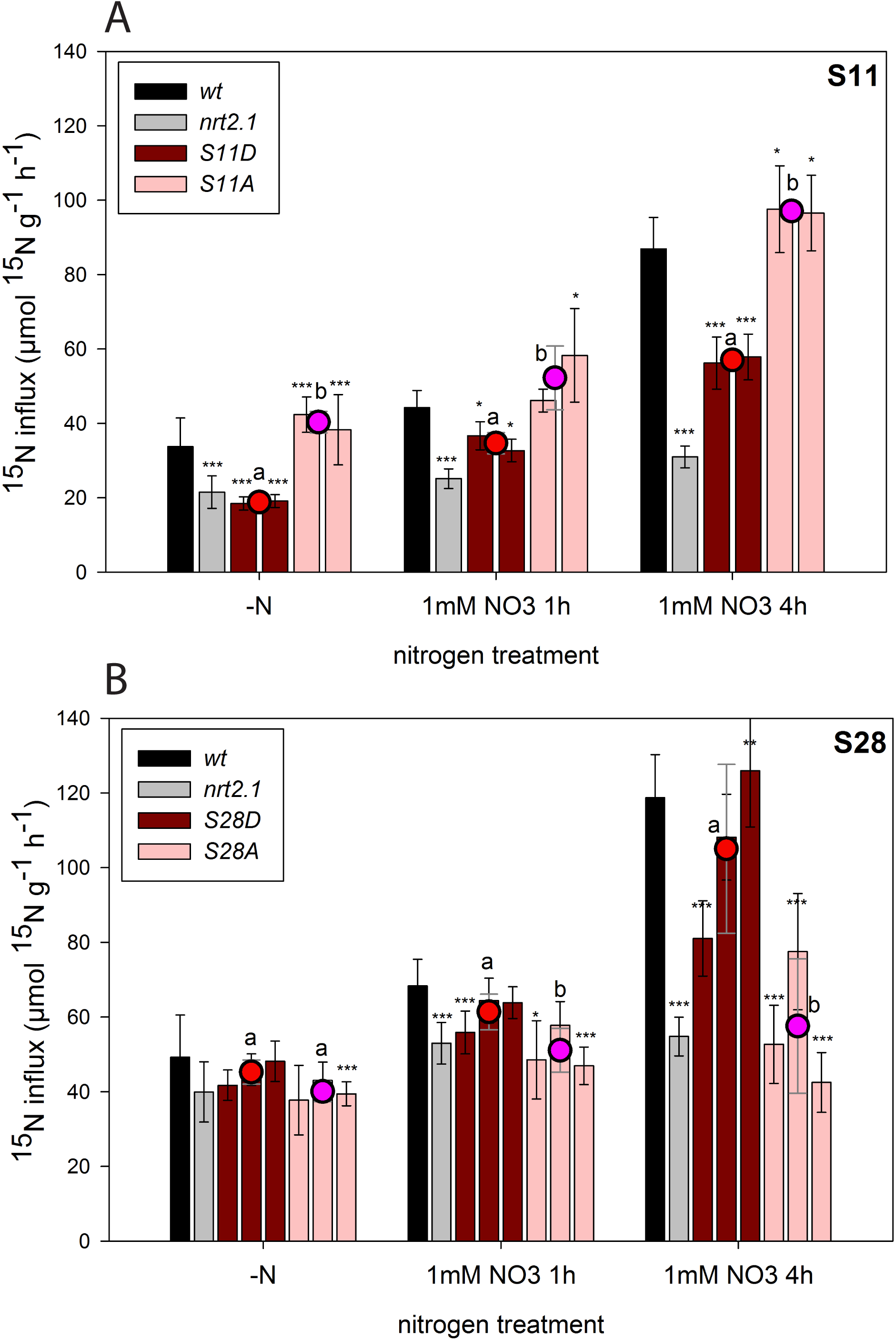
Regulation of nitrate uptake by N-terminal NRT2.1 phosphorylation at (**A**) S11, and (**B**) S28. In all panels, wild type and *nrt2.1* knock out mutant were used as controls. Phosphorylation site mutants of *NRT2.1* were expressed under the *NRT2.1* promoter in the *nrt2.1* mutant background as phosphomimicking (S to D) or phosphodead (S to A) version of different phosphorylation sites. Bars of same color indicate independent lines expressing the same construct (S to D, or S to A). Averages with standard deviation are displayed, asterisks indicate significant differences to wild type within one treatment (*** p<0.001, ** p<0.01, * p<0.05, pairwise t-test). Dots indicate average uptake rates of different plant lines expressing S to D or S to A mutants. Letters indicate significant differences (p<0.05, pairwise t-test) between phosphomimicking and phosphodead mutants within each treatment.

### Phosphorylation-dependent NRT2.1 regulation through interaction with NAR2.1

Nitrate transporter NRT2.1 was previously shown to be regulated by the interaction with protein NAR2.1 (AT5G50200) (Laugier et al., 2012), and this interaction resulted in activation of NRT2.1 transport activity (Orsel et al., 2006; Orsel et al., 2007). Therefore, we used ratiometric bimolecular fluorescence complementation (Grefen and Blatt, 2012) to explore whether the interaction of NRT2.1 with NAR2.1 was affected by NRT2.1 phosphorylation (Figure 2A). The known interaction of NRT2.1 with NAR2.1 was used as a reference. Putative phosphomimic mutation (S11D) at S11 did not result in differential interaction with NAR2.1 compared to phosphodead mutations (S11A), but in both cases the interaction of NAR2.1 was stronger than with non-mutated wild type NRT2.1. Interestingly, for S21 we observed a significantly increased interaction of phosphodead NRT2.1 S21A with NAR2.1 compared to phosphomimicking NRT2.1S21D, which displayed a weaker interaction with NAR2.1. By contrast, NRT2.1 with phosphodead mutation S28A showed significantly low interaction with NAR2.1, while phosphomimicking NRT2.1S28D displayed stronger interaction with NAR2.1. Thus, phosphorylation at S21 and S28 affected the interaction of NRT2.1 with its activator NAR2.1 in opposite ways: The interaction of NRT2.1 with NAR2.1 was favored by phosphorylation at S28, and was less favored by phosphorylation at S21 (Figure 2B, C).

**Figure 2:**
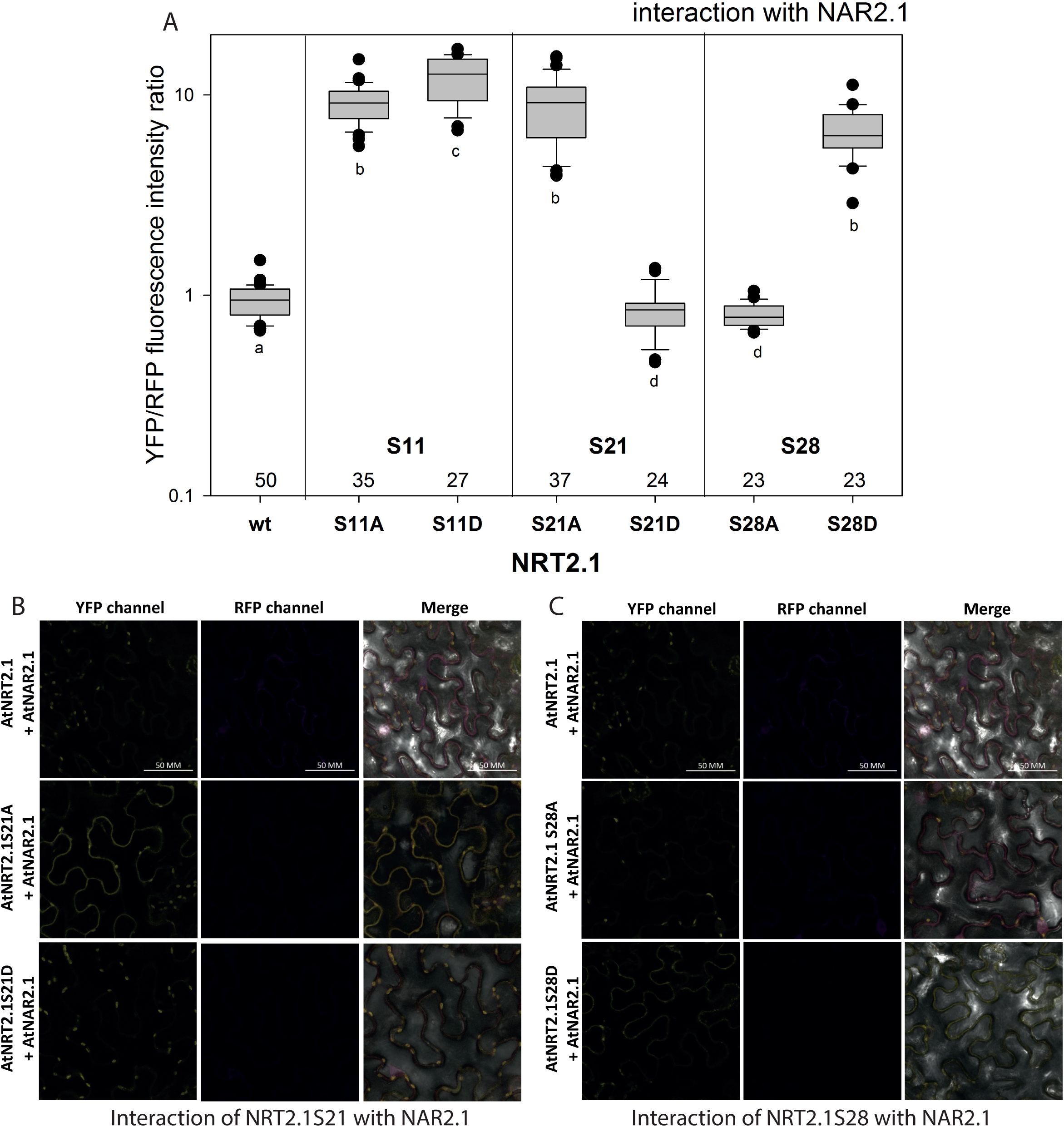
Interaction of NRT2.1 with NAR2.1. (**A**) The effect of phosphodead (S to A) and phosphomimicking (S to D) mutations of NRT2.1 phosphorylation sites on the interaction with NAR2.1. Numbers indicate the total number of cells quantified. Small letters indicates significant differences as determined by pairwise rank sum test (p<0.05) (**B**) Representative images of the interaction of NRT2.1S21A and NRT2.1S21D with NAR2.1 (YPF-channel) and plasmid expression control (RFP-channel) (**C**) Representative images of the interaction of NRT2.1S28A and NRT2.1S28D with NAR2.1 (YPF-channel) and plasmid expression control (RFP-channel). Scale bar: 50 µm.

### Identification of kinases phosphorylating NRT2.1

Based on existing nitrate starvation and nitrate resupply data sets (Engelsberger and Schulze, 2012; Menz et al., 2016), and under consideration of the newly performed resupply-experiment with 0.2mM and 5 mM nitrate, we aimed to identify kinases phosphorylating the different phosphorylation sites of NRT2.1. We hypothesize that a kinase phosphorylating NRT2.1 is also likely to be regulated by nitrate-dependent phosphorylation. Thus, phosphorylation profiles of the kinase and its substrate should be highly correlated (if phosphorylations at kinase and substrate both have activating, or both have inactivating effects) or anti-correlated (if one activating and one inactivating phosphorylation occurs). To identify such kinase candidates, phosphorylation profiles of NRT2.1 phosphorylation sites S11, S28, S501 and T521 were correlated with the phosphorylation profiles of phosphorylation sites in identified kinases (Supplementary Figure S3, Supplementary Table S2). NRT2.1 phosphorylation site S21 was not included due to low data coverage across conditions. Following this approach, phosphorylation profile correlation pairs were strictly filtered based on their correlation r-value (r > 0.9 or r < −0.9). As a result, 12 protein kinases were identified under nitrate deprivation, 34 protein kinases were identified only under nitrate resupply conditions (5 mM NO_3_ or 0.2 mM NO_3_) and 15 protein kinase were identified under nitrate depletion as well as under nitrate resupply (Figure 3A, Supplementary Table 2).

**Figure 3:**
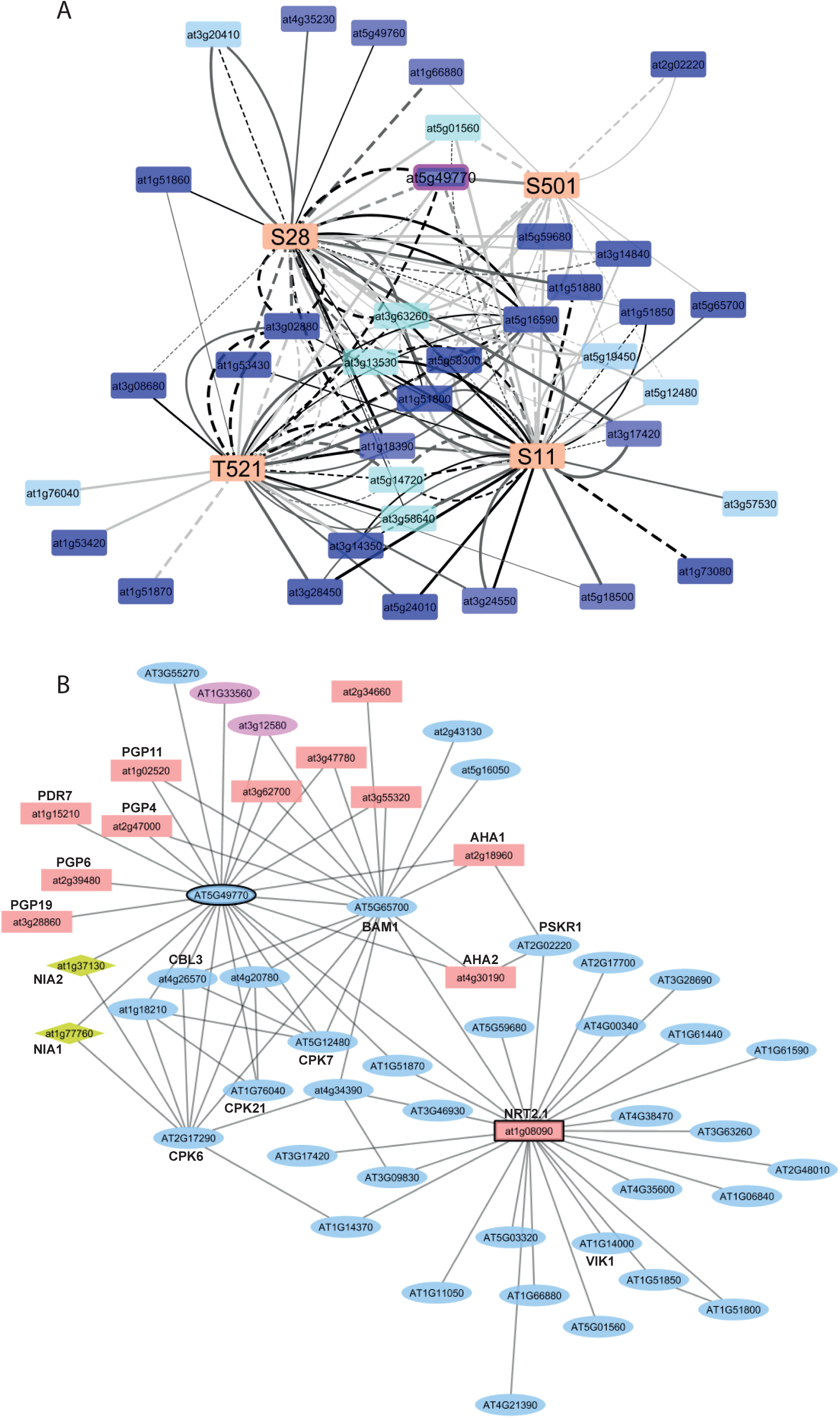
Phosphorylation and interaction network of NRT2.1 and its putative kinases (**A**) Correlation network of NRT2.1 phosphorylation sites S11, S28, S501 and T521 with plasma membrane located kinases under nitrate deprivation and nitrate resupply with 5mM and 0.2 mM NO3. Solid edges: positive correlation, dashed edges: negative correlation. Edge color: black, 5mM NO3; gray, 0.2mM NO3; light gray, nitrogen deprivation. Node color: blue, receptor kinases; light blue, calcium signaling; gray, other kinases. (**B**) Interaction network of NRT2.1 and the selected putative kinase AT5G49770 extended by known interaction partners from STRING (Franceschini et al., 2013). Node color: red, transport proteins; blue, signaling proteins; yellow, nitrogen metabolism.

Among the kinases which displayed phosphorylation profiles highly correlated with NRT2.1 phosphorylation, we found MAPKKK7 (AT3G13530), for which its own phosphorylation positively correlated with phosphorylation of NRT2.1 at with S11 and T521. Phosphorylation of CALCIUM-DEPENDENT PROTEIN KINASE 29 (CPK29, AT1G76040) was highly correlated with NRT2.1 T521 phosphorylation, while phosphorylation of CPK19 (AT5G19450) and CPK7 (AT5G14720) at different sites was connected to the phosphorylation profiles of NRT2.1 at S11, S28, and S501. Among the receptor kinases, phosphorylation of BAM1 (AT5G65700) positively correlated with S11 and S501 phosphorylation, and phosphorylation of PHYTOSULFOKINE RECEPTOR 1 (PSKR1, AT2G02220) correlated with phosphorylation at S501. C-terminal phosphorylation at LIEEVSHSSGS(ph)PNPVS(ph)D of co-receptor QSK1 (AG3G02880) showed a negative correlation with phosphorylation at S28 and T521. Phosphorylation of kinase AT5G49770 at S839 showed positive correlation with NRT2.1 phosphorylation at S28 under low nitrate supply, and negative correlation with S28 phosphorylation at high nitrate supply (Supplementary Table S2). Receptor kinase candidate AT5G49770 raised our further interest, since it was found to be multiply phosphorylated. Phosphorylation sites were identified within the activation loop at T792 (LVGDPEKAHVT(ph)TQVK), in the kinase domain at S839 (SPIDRGS(ph)YVVK) and S870 (NLYDLQELLDTTIIQNS(ph)GNLKGFEK), and within the disordered C-terminal region at S919 (LVGLNPNADS(ph)ATYEEASGDPYGR) (Supplementary Figure S4).

The phosphorylation network was then further complemented with known interaction partners of NRT2.1 and kinase candidate AT5G49770 as retrieved from public sources (Franceschini et al., 2013) (Figure 3B). Thereby, it became obvious that NRT2.1 interacted with several other kinases besides AT5G49770, among them VIK1 (AT1G14000) (Wingenter et al., 2011), a MAP kinase (AT3G46930), phytosulfokine receptor kinase PSKR1 (AT2G02220), and several uncharacterized receptor kinases (Figure 3B). By contrast, kinase AT5G49770 was found to interact with various other transport proteins within the membrane besides NRT2.1, for example with plasma membrane ATPases AHA1 (AT2G18960) and AHA2 (AT4G30190), and several members of the ABC transporter family (PGP6 AT2G39480, PGP19 AT3G28860, PDR7 AT1G15210, PGP4 AT2G47000, PGP11 AT1G02520, MRP10 AT3G62700, PGP20 AT3G55320). Furthermore, kinase AT5G49770 connected to the calcium signaling pathway through interactions with CPK29 (AT1G76040), CPK7 (AT5G12480), CPK6 (AT2G17290), and CBL3 (AT4G26570). A connection to nitrate assimilation was found trough the interaction of AT5G49770 with nitrate reductases NIA2 (AT1G37130) and NIA1 (AT1G77760).

### Kinase AT5G49770 interacts with and phosphorylates NRT2.1

Kinase candidate AT5G49770 was then tested for its ability to in planta interact with NRT2.1 and we further tested whether this interaction was affected by the phosphorylation of the kinase at its four phosphorylation sites. Indeed, we could show an interaction of AT5G49770 with NRT2.1 in the ratiometric bimolecular fluorescence complementation system (Figure 4A, Supplementary Figure S5A). Phosphorylation status of AT5G49770 at sites T792, S870 and S919 did not differentially affect the interaction with NRT2.1. However, if S839 was mutated to a phosphodead alanine, the interaction of AT5G49770S839A with NRT2.1 was significantly increased compared to AT5G49770 in which S839 was mutated to phosphomimicking aspartate. We conclude that AT5G49770 mimicking dephosphorylated S839 (S839A) showed stronger interaction with NRT2.1 than AT5G49770 mimicking a phosphorylated state (S839D). Next, we explored, if NRT2.1 actually is a substrate for kinase AT5G49770 (Figure 4B). The intracellular domain of AT5G49770 was recombinant expressed and exposed to the substrate peptide EQSFAFSVQSPIVHTDK in in-vitro kinase assays (Wu and Schulze, 2015). Indeed, AT5G49770 was able to phosphorylate the substrate peptide at S21. Significantly higher S21 phosphorylation was observed in recombinant kinase domains with phosphodead mutation AT5G49770S839A. By contrast, phosphomimicking mutation AT5G49770S839D resulted in lower substrate phosphorylation at S21. Also for phosphorylation site S870, phosphorylation site mutations resulted in differential substrate phosphorylation efficiency: phosphomimicking mutation AT5G49770S870D resulted in significantly higher kinase activity towards NRT2.1 N-terminal peptide than phosphodead mutation AT5G49770S870A. No significant difference in substrate phosphorylation between phosphomimicking and phosphodead versions of recombinant AT5G49770 kinase domain were observed for other phosphorylation sites of the kinase (Figure 4B). Consistently, in all *in vitro* kinase activity assays, phosphorylation of the substrate peptide was only found at the serine corresponding to NRT2.1S21 (Figure 4C), and never at the site corresponding to S28 or the doubly phosphorylated forms.

**Figure 4:**
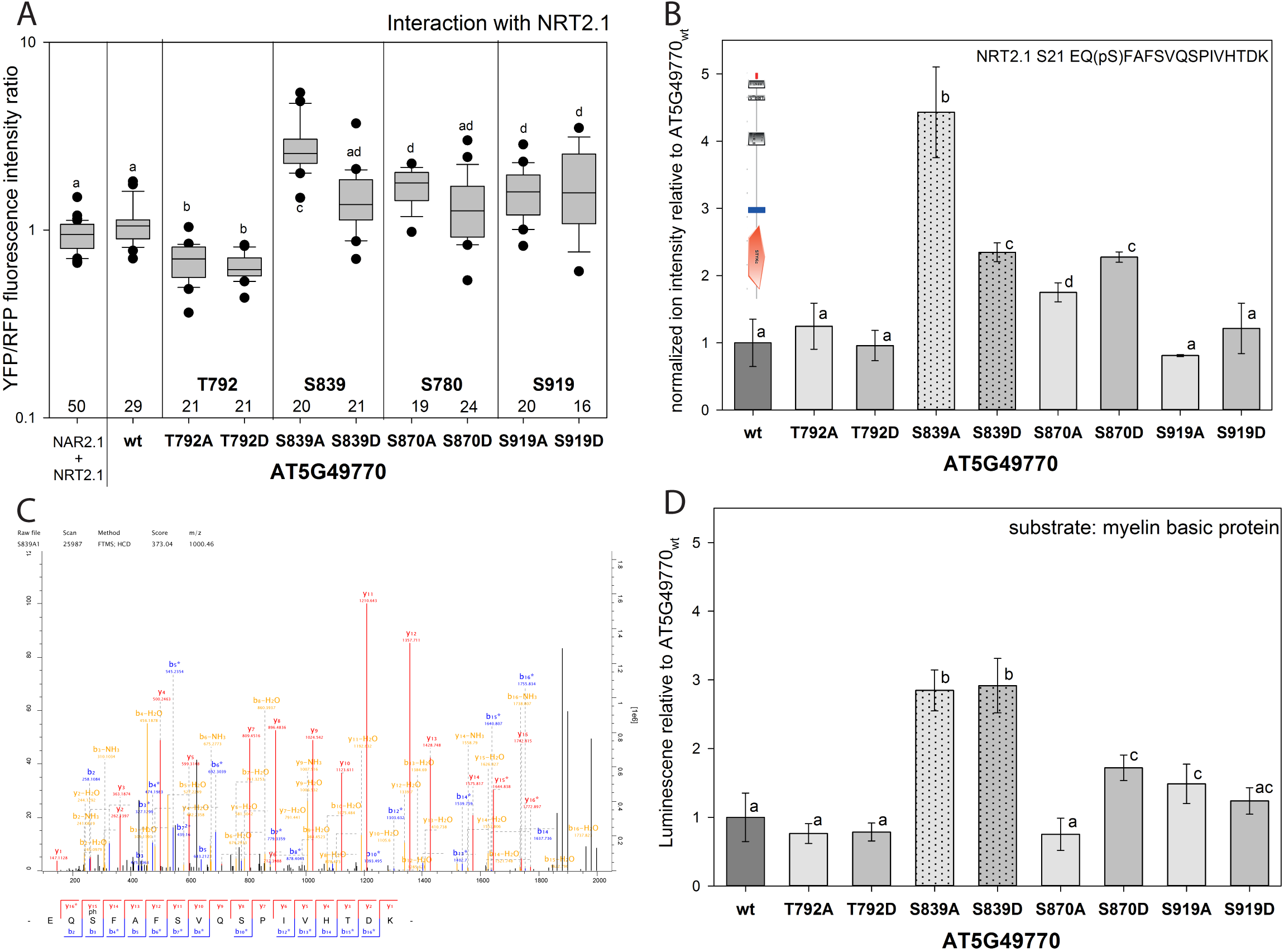
Kinase AT5G49770 interacts with and phosphorylates NRT2.1 at S21. (A) The effect of phosphodead (S to A) and phosphomimicking (S to D) mutations of AT5G49770 phosphorylation sites on the interaction with NRT2.1. Numbers indicate the total number of cells quantified. Small letters indicates significant differences as determined by pairwise rank sum test (p<0.05). (**B**) *In vitro* kinase activity assay using the NRT2.1 peptide EQSFAFSVQSPIVHTDK as a substrate. Averages with standard deviations are displayed, small letters indicates significant differences based on pairwise t-test (p<0.05). Dotted bars denote phosphorylation site S839. Domain model of the AT5G49770 kinase based on SMART is displayed as insert (Schultz et al., 1998). (**C**) Representative fragment spectrum of the *in vitro* kinase activity assay showing specific phosphorylation of the substrate peptide EQSFAFSVQSPIVHTDK at S21. Spectrum was directly exported from MaxQuant version 1.5.3.30 (Cox and Mann, 2008). (**D**) *In vitro* kinase activity assay using myelin basic protein as a substrate. Averages with standard deviations are displayed, small letters indicates significant differences based on pairwise t-test (p<0.05).

Interestingly, when the generic kinase substrate myelin basic protein was used in the in-vitro kinase assay instead of the specific NRT2.1 substrate peptide, phosphorylation site mutations at AT5G49770 S839 did not result in differential kinase activity, and increased kinase activity was only observed for AT5G49770S870D mutation (Figure 4D). These results suggest that phosphorylation of AT5G49770 at S870 regulated general kinase activity (i.e. also towards a generic substrate), while phosphorylation of AT5G49770 at S839 rather affected the interaction with the kinase substrate: When AT5G49770 S839 was phosphorylated (i.e. phosphomimicked in S839D), interaction of AT5G49770 with NRT2.1 was reduced (Figure 4A) and lower levels of substrate phosphorylation at S21 occurred (Figure 4B). When S870 was phosphorylated (i.e. phosphomimicked in S870D), kinase activity generally was slightly enhanced (Figure 4D), but this did not affect the interaction of AT5G49770 with NRT2.1 (Figure 4A).

We then checked the *in-planta* phosphorylation status of NRT2.1 and AT5G49770 (Figure 5) in the two nitrogen nutrition experiments, such as a nitrogen deprivation experiment with nitrate deprivation for 15 minutes and 3 hours (Menz et al., 2016), and a nitrate starvation resupply experiment with two days of nitrate starvation followed by resupply of either 0.2 mM or 5 mM nitrate for 5 minutes or 15 minutes. A trend of NRT2.1 S21 to be increasingly phosphorylated under nitrate resupply conditions with high (5mM) nitrate was observed. No phosphorylation was identified under prolonged nitrate starvation for 2 days (Figure 5A). By contrast, NRT2.1 S28 was strongly phosphorylated under nitrate starvation and showed a trend to be dephosphorylated with resupply of high nitrate concentrations (5 mM) (Figure 5B). This confirms earlier observations of NRT2.1 S28 dephosphorylation upon nitrate resupply (Engelsberger and Schulze, 2012). Kinase AT5G49770 was found with increasing phosphorylation at S870 after resupply of 5 mM nitrate (Figure 5C) and we observed dephosphorylation trends at S839 under high (5 mM) nitrate resupply (Figure 5D). The kinase phosphorylation sites were only detected at very low levels under nitrogen starvation and resupply with low (0.2 mM) nitrate concentrations. In summary, NRT2.1 phosphorylation at S21 and S28 showed a tendency for anti-correlation across all conditions: When S21 was highly phosphorylated, phosphorylation at S28 was detected at lower level and *vice versa*.

**Figure 5:**
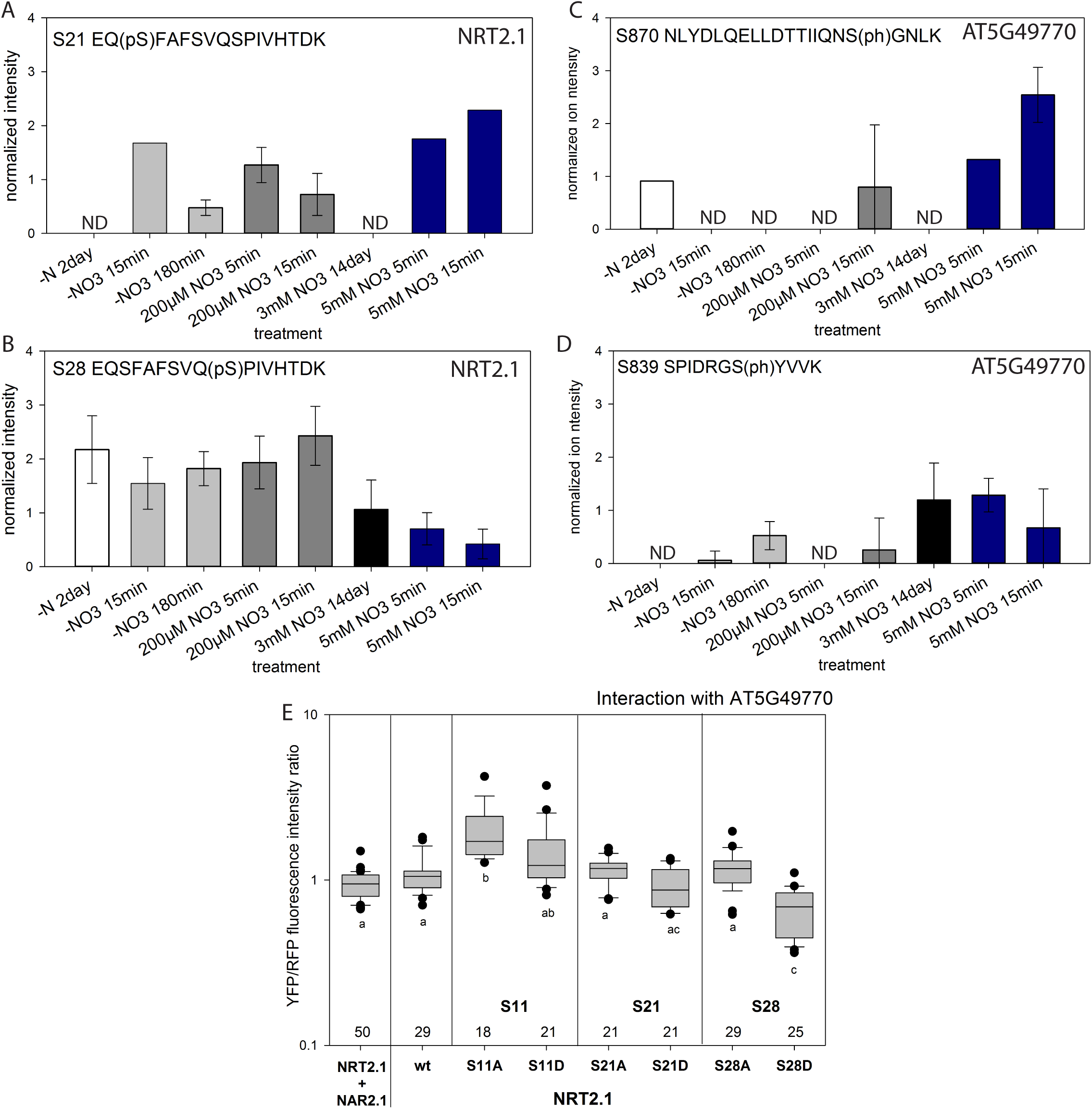
Phosphorylation status of NRT2.1 S21, S28 and AT5G49770 S870, S839 under different nutritional context. (**A**) Phosphorylation of NRT2.1 at S21. (**B**) Phosphorylation of NRT2.1 at S28. (**C**) Phosphorylation of AT5G49770 at S870, (D) Phosphorylation of AT5G49770 at S839. In panels (A) to (D) raw files from previous experiments (Engelsberger and Schulze, 2012; Menz et al., 2016) and current experiment of resupply with 0.2 mM and 5 mM nitrate were processed together and quantified. Average values of three biological replicates are shown with standard deviation as determined by cRacker (Zauber and Schulze, 2012). Averages of three replicates are shown with standard deviations. ND: not detected. (**E**) Interaction of AT5G49770 with NRT2.1 and respective NRT2.1 phosphorylation site mutants as determined from ratiometric bimolecular fluorescence experiments. Numbers indicate the number of quantified cells.

### Kinase AT5G49770 affects nitrate uptake

Interestingly, the interaction of kinase AT5G49770 with NRT2.1 was found to be affected by the NRT2.1 phosphorylation status (Figure 6A, Supplementary Figure S5B): AT5G49770 showed significantly weaker interactions with NRT2.1 in the phosphorylation mimicking NRT2.1S28D mutant compared to phosphodead NRT2.1S28A. This suggests that the interaction of the kinase with NRT2.1 is prevented by phosphorylation of NRT2.1 at site S28 (Figure 5E). The interaction of the kinase with NRT2.1 remained largely unaffected by the phosphorylation status of other N-terminal phosphorylation sites S21 and S11. Since NRT2.1 activity was lower when S28 was dephosphorylated (i.e. phosphodead NRT2.1S28A) compared to when S28 was phosphorylated (i.e. phosphomimicking NRT2.1S28D) (Figure 1), kinase AT5G49770 seems to preferentially interact with NRT2.1 in its inactive state.

**Figure 6:**
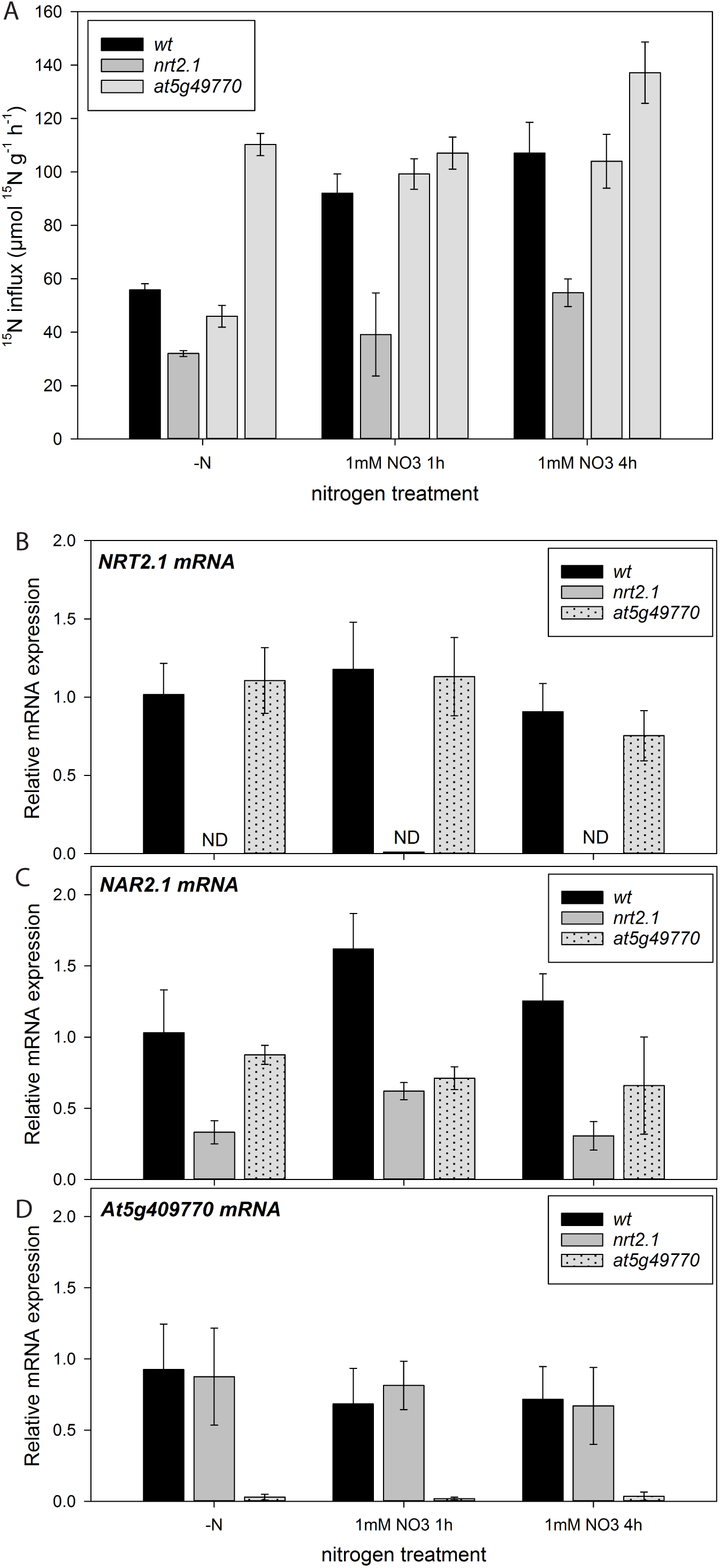
Interaction of AT5G49770 with NRT2.1 phosphorylation site mutants and nitrate uptake of kinase knock-out. (**A**) Nitrate influx rate of wild type, *nrt2.1* knock-out and *at5g49770* knock-out under nitrate starvation (-N) and induction with 1 mM nitrate for 1hour and 4hours. Averages with standard deviations are shown. (**B**) Expression of *NRT2.1* (**C**) *NAR2.1* mRNA and (**D**) *At5g49770* mRNA in wild type, *nrt2.1* knock-out and *at5g49770* knock-out mutant. Averages of three biological replicates are shown with standard deviation.

Consequently, in the knock-out *at5G49770* we observed a higher nitrate influx than in *nrt2.1-2* knock out (Figure 6A), reaching wild type like or even higher than wild type nitrate influx rates under nitrate induction (1 mM NO_3_ for 1h and 4h). Thus, in absence of AT5G49770, the nitrate influx was high, consistent with high activity of NRT2.1. We therefore propose that kinase AT5G49770 acts as an inhibitor to NRT2.1 transporter, and therefore we propose to name the kinase as NITRATE UPTAKE REGULATORY KINASE NURK1. The observed changes in the nitrate uptake rate in *nurk1* (*at5g40770*) knock-out mutant were not connected to altered *NRT2.1* mRNA levels (Figure 6B). However, a slightly reduced expression of *NAR2.1* mRNA was observed in the kinase mutant *at5g49770* (Figure 6C), suggesting a possibility for downstream signaling effects. Expression of the kinase was not dependent on the presence of NRT2.1, since *nrt2.1-2* mutant showed similar levels of *Nurk1* (*At5g49770*) mRNA as wild type, at least under the conditions tested here (Figure 6D).

### Functional model of the S21-S28 phospho-switch

Based the results above, the N-terminus of NRT2.1 was found to undergo different interactions, either with NAR2.1 or kinase NURK1, dependent on the phosphorylation status of the two serines at S21/S28. Also receptor kinase NURK1 was found to be regulated by two phosphorylation sites, which affect kinase activity and the interaction with NRT2.1. Thus, NRT2.1 and NURK1 can switch between active and inactive states depending on their phosphorylation patterns:

Under conditions when NRT2.1 transport is active, preferentially at low external nitrate supply or under nitrate starvation, NRT2.1 is phosphorylated at S28 and interaction with NAR2.1 can take place to stabilize this active state (Figure 7 “yang”). At the same time, when NURK1 is phosphorylated at S839 it does not interact with NRT2.1, Most likely the kinase under these conditions is also dephosphorylated at S870, corresponding to an inactive kinase domain. Under conditions when NRT2.1 transport is inactive, NRT2.1 is dephosphorylated at S28. The dephosphorylated state of NRT2.1 enables the interaction with receptor kinase NURK1. The kinase in turn can then phosphorylate NRT2.1 at S21 to stabilize the inactive state of NRT2.1 by preventing interactions of NRT2.1 with NAR2.1 (Figure 7 “yin”). The interaction of NURK1 with NRT2.1 requires (i) dephosphorylation of NRT2.1 S28 by phosphatases, and (ii) dephosphorylation of the kinase NURK1 at S839, and (iii) concurrent activation of the kinase by (auto)phosphorylation at S870. In turn, switching NRT2.1 from inactive state to active transport requires de-phosphorylation at S21 by a phosphatase and, more importantly, phosphorylation at S28 by another kinase.

**Figure 7:**
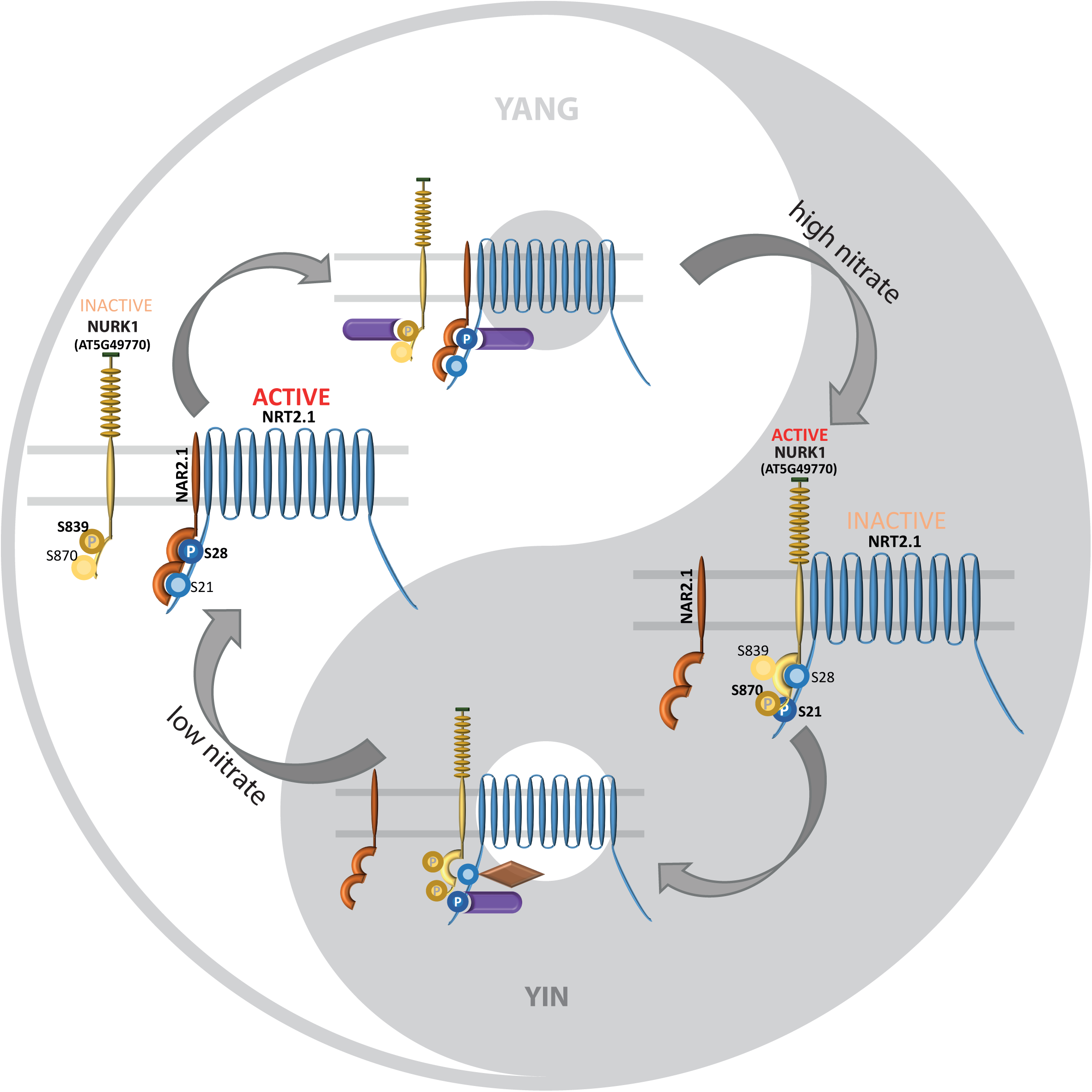
Functional model of two contrasting activity states of NRT2.1 are stabilized by contrasting interactions of NRT2.1 with either NAR2.1 or NURK1 (AT5G49770). **Yang:** Active state of NRT2.1 is characterized by phosphorylation at S28 and interaction with NAR2.1. Kinase NURK1 does not interact with NRT2.1 and is phosphorylated at S839. **Yin:** Activation of kinase NURK1 is achieved by dephosphorylation at S389 and phosphorylation at S870. NURK1 can phosphorylate NRT2.1 at S21, thereby preventing the interaction of NRT2.1 with NAR2.1. Functional relevance of other NRT2.1 phosphorylation sites was not considered in this figure. Purple: phosphatases, brown: another kinase.

Possibly structural effects favor the interaction of NRT2.1 with its activator NAR2.1 only when S28 is phosphorylated, while the interaction of NRT2.1 with the kinase is favored under conditions when S28 is not phosphorylated. In turn, phosphorylation at S21 prevents the interaction of NRT2.1 with NAR2.1. The switch of phosphorylation state of NRT2.1 between S21 and S28 must happen very rapidly, since a doubly-phosphorylated peptide was only rarely detected when nitrate was resupplied to starved plants at 0.2 mM and 5 mM (Supplementary Figure S6). The functional role of receptor kinase NURK1 thus can be hypothesized in switching to and stabilization of inactive state of NRT2.1. Thereby, NURK1 itself seems to be activated by (auto)phosphorylation at S870, and dephosphorylation at S839 within its kinase domain enhances the ability to interact with NRT2.1 at de-phosphorylated S28.

Thus, serines S21 and S28 in the N-terminus of NRT2.1 act like a yin-yang phosphoswitch by which the activating interaction of NRT2.1 with NAR2.1 is enabled or disabled. The interaction of the kinase NURK1 with NRT2.1 – which replaces the interaction of NAR2.1 with NRT2.1 – is further fine-tuned by another phosphoswitch within the kinase at serines S839 and S870. Activity changes of NRT2.1 and resulting effects on nitrate transport are a final result. We postulate, that the phosphoswitch mechanism requires fast action also of yet unknown phosphatases, which act both on NURK1 and NRT2.1. Furthermore, action of another kinase is required to phosphorylate S28 upon switching NRT2.1 from inactive to active state (Figure 7).

## Discussion

Regulation of NRT2.1 is very complex. There are (at least) five phosphorylation sites in the N- and C-terminus which differentially affect nitrate uptake. In the C-terminus, phosphorylation of S501 was recently shown to result in inactivation of NRT2.1 by favoring a multimerization state with closed transport pores in contrast to monomeric state with open pores at S501 dephosphorylation (https://www.biorxiv.org/content/10.1101/583542v1). In the N-terminus, phosphorylation (or phosphomimic) of S28 mediate high nitrate uptake, while phosphorylation at S11 resulted in low nitrate uptake. Thus, NRT2.1 seems to be activated by phosphorylation at S28 and inhibited by phosphorylation at S11. The molecular mechanisms by which this posttranslational control at these different sites takes effect, can be very different for each of these sites. Here, we discovered a phospho-switch mechanism in the NRT2.1 N-terminus involving S21 and S28, and by which the interaction of NRT2.1 with its activator NAR2.1 is affected through competition with the interaction of NURK1 at the same NRT2.1 phosphorylation site S28. A role of phosphorylation of NRT2.1 at S28 has recently also been proposed to affect protein stability (Zou et al., 2019).

It is well known that phosphorylation of the same protein at different sites, or the existence of different modifications at closely neighboring sites can influence the nature of protein-protein interactions and consequently the activity of proteins or signaling pathways. Classic examples are mammalian tyrosine receptor kinases, in which multiple tyrosine phosphorylation sites recruit different interaction partners to activate distinct downstream signaling cascades (Schulze et al., 2005). More recently, multisite phosphorylation in ribosomal protein S6 kinase 1 (S6K1) was shown to alter substrate specificity by the kinase (Arif et al., 2018). Thereby, phosphorylation of three different sites was required for S6K1 to phosphorylate one substrate but not the other. In receptor kinase signaling, a tyrosine phosphorylation switch has recently been discovered in SERK3/BAK1 which in the phosphorylated state activates immune signaling, and in the dephosphorylated state feeds into other signaling pathways (Perraki et al., 2018). Modification-dependent activity regulation at two neighboring sites can also occur by two different modifications: A phospho/metyl-switch at histone H3 was shown to regulate TFIID association with the chromosomes (Varier et al., 2010). High affinity binding of the transcription factor to the respective promoter sequence was enabled by histone 3 trimethylation, but inhibited by phosphorylation of histone 3 at an adjacent threonine. In analogy, serine phosphorylation and oxidation of a nearby methionine had antagonistic effects on the activity of sucrose-phosphate synthase (Hardin et al., 2009). Here, we present an example in which phosphorylation at spatially closely neighboring phosphorylation sites in the N-terminus of NRT2.1 resulted in recruitment of different interaction partners. This differential recruitment of interacting proteins as a final consequence resulted in active or inactive states of the major nitrate uptake transporter NRT2.1.

NITRATE UPTAKE REGULATORY KINASE NURK1 is highly and almost exclusively expressed in root tissue, particularly in the lateral root cap (Winter et al., 2007) and shows high overlap with NRT2.1 expression. Within the Arabidopsis proteome there are two closely related kinases (AT5G49760 and AT5G49780) sharing high homology to our candidate kinase NURK1, AT5G49770 (Supplementary Figure S7). AT5G49760 was highly expressed in leaf tissue and anthers, and showed rather low expression values in roots, and AT5G49680 showed moderate general expression also in roots, but at lower levels than NURK1 (Winter et al., 2007). All phosphopeptides identified for NURK1 were proteotypic peptides clearly identifying the AT5G49770 protein. Interestingly, all four phosphorylation sites identified for NURK1 except S839 are highly conserved across all three homologs. S839 is unique to AT5G49770, while in AT5G49760 and AT5G49780 a lysine is present at the respective position within the sequence (Supplementary Figure S7). In the kinase activity assays, AT5G497780 was largely inactive, while AT5G497760 displayed a higher activity towards substrate peptide EQSFAFSVQSPIVHTDK than did AT5G49770. This high activity could possibly be explained by the fact that the regulatory phosphorylation site identified for NURK1 with S839 is not present in AT5G49760 and therefore this kinase may show constitutively higher activity. Since the *in vitro* phosphorylation assay was conducted with recombinant kinase domains of each kinase homolog, the native kinase domain of NURK1 was expected not to be highly phosphorylated and thus yielded lower activity than recombinant kinase domain with phosphomimicking S839D mutations. In fact, the kinase activity of AT5G49770S839D was similarly high as the activity of AT4G49760 (Figure 4). We cannot provide clear explanation for the observed inactivity of recombinant kinase domain AT5G49780, possibly this is connected to the additional sequence at amino acids 609 to 629 (Supplementary Figure S7).

Protein kinases can be regulated by phosphorylation within their kinase domain. Most prominent examples are the mitogen-activated protein kinases (MAP-Kinases), which are activated by double-phosphorylation of a well-conserved TEY motif within the activation loop (Rodriguez et al., 2010). Also in SUCROSE NON-FERMENTING 1-RELATED PROTEIN KINASE 1 (SnRK1), phosphorylation of the kinase domain is required for full activity (Emanuelle et al., 2018). Kinase candidate NURK1 was found to have four phosphorylation sites within its kinase domain. The phosphorylation status of S839 and S879 resulted in differential activity towards the substrate NRT2.1 when mutated to phosphomimicking or phosphodead amino acids. Thereby, dephosphorylation at S839 (phosphodead mutation) was associated with higher kinase activity and stronger interaction with the substrate. Interestingly, phosphomimicking mutation at S870 also increased kinase activity towards generic substrate myelin basic protein, while mutations at S839 did not alter kinase activity towards the generic substrate. This suggests that S870 is a phosphorylation site regulating general kinase activity, while phosphorylation at S839 affects the specific interaction with the substrate protein NRT2.1. Phosphorylation site T792 was predicted (InterPro, (Mitchell et al., 2019)) to be in the activation loop of the kinase. In contrast to other well-characterized kinases, phosphomimicking or phosphodead mutations of T792 within the activation loop did not affect the activity of kinase NURK1. There are several kinases in which phosphorylation of the activation loop is not required for activation (Nolen et al., 2004). Possibly, in NURK1, steric access to the activation loop and the active center is regulated by (de)phosphorylation of S839 rather than by phosphorylation of the activation loop. Recent structural analyses of kinase catalytic domains revealed that the activation loop requires N- and C-terminal structural anchors which correctly place the activation loop neighboring the ATP-binding site and catalytic domain (Nolen et al., 2004). Based on this model, in NURK1 phosphorylation site S839 is located within a region that could serve as C-terminal anchor for the activation loop. However, the precise role of the different phosphorylation sites of NURK1 need to be explored in further experiments.

Regulation of transporters at the plasma membrane by multiple phosphorylation sites could well be a general principle, particularly for proteins that are under control of many different signals. For example, the plasma membrane ATPase AHA2 is regulated by two different activating phosphorylation sites in its C-terminus and by at least one inactivating phosphorylation site in one of the cytosolic loops. In addition, these regulatory phosphorylation sites of AHA2 are targeted by different kinases which control proton flux in response to different stimuli (Fuglsang et al., 2003; Haruta et al., 2008; Caesar et al., 2011; Fuglsang et al., 2014; Haruta et al., 2015). Also for aquaporins, different regulatory modifications were described with different effects on pore gating and/or membrane localization (Tornroth-Horsefield et al., 2006; Prak et al., 2008; di Pietro et al., 2013), and a kinase was identified to regulate aquaporins in response to sucrose (Wu et al., 2013). NRT2.1 here was found to be multiphosphorylated in the N- and C-terminus. At the same time, the regulatory pattern of NRT2.1 turned out to be very complex and under transcriptional and posttranscriptional control of external nitrate supply as well as the metabolic C/N status of the plant (Gansel et al., 2001; Wirth et al., 2007; Camanes et al., 2012; Li et al., 2016). Therefore, it is to no surprise at all to find highly complex regulatory mechanisms involving a yin-yang like phosphoswitch to control NRT2.1 activity at the protein level.

## Conclusions

This work functionally characterizes two phosphorylation sites which work together as a phospho-switch in the N-terminus of NRT2.1. Thereby, NRT2.1 is not regulated directly, but differential phosphorylation at S28 and S21 modulates the interaction of NRT2.1 with the activator protein NAR2.1. Since the interaction of NRT2.1 with NAR2.1 is essential for NRT2.1 activity (Laugier et al., 2012), the S21/S28 phospho-switch is likely to be a major regulator of nitrate uptake into root cells through NRT2.1. We propose NURK1 as a kinase that controls S21 phosphorylation and thus inhibits the interaction of NRT2.1 with activator NAR2.1 leading to low activity rates of NRT2.1, especially under high nitrate resupply.

## Supporting information

Supplementary Material Description

## Acknowledgements

We thank Zhaoxia Zhang for help with bioinformatics scripting and Sven Gombos for technical assistance in the lab. The work was supported by an international grant from the ANR in France (SIPHON ANR-13-ISV6-0002-01) and Deutsche Forschungsgemeinschaft (SCHU1533/8-1) in Germany.

