## Supplementary figures and images for "A phospho-switch in the N-terminus of NRT2.1 affects nitrate uptake by controlling the interaction of NRT2.1 with NAR2.1"

### Supplementary Figure S1

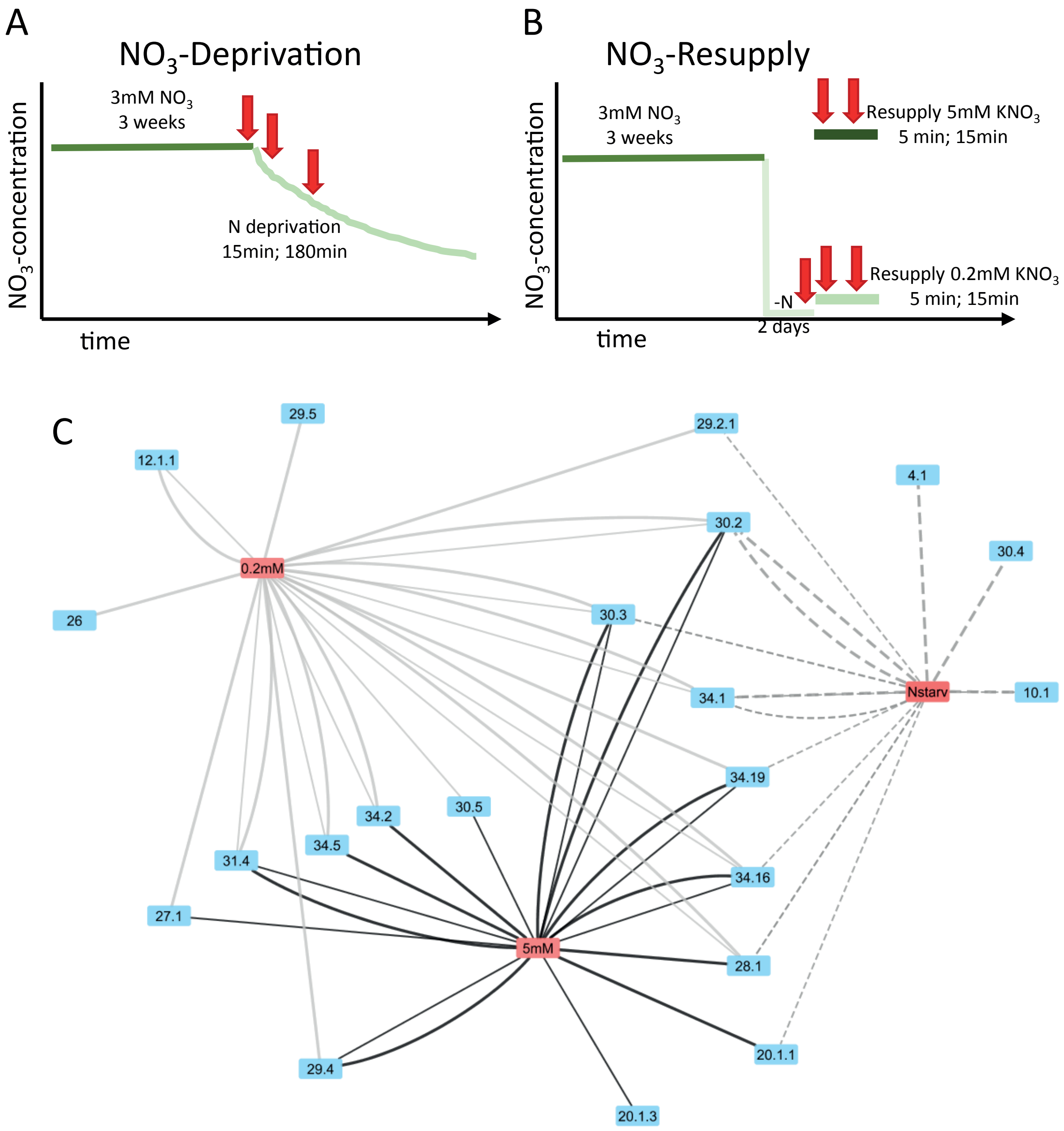

### Supplementary Figure S2

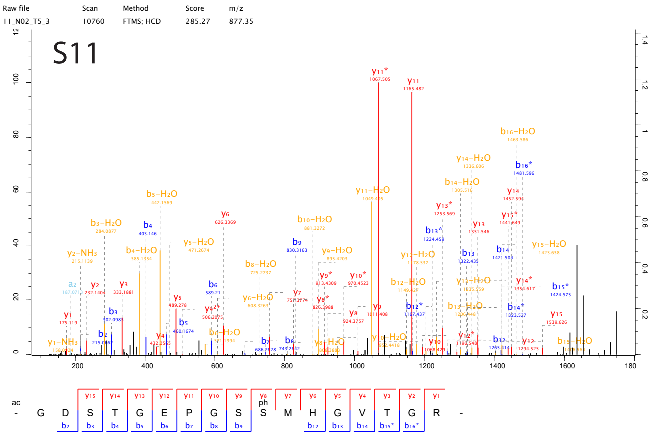

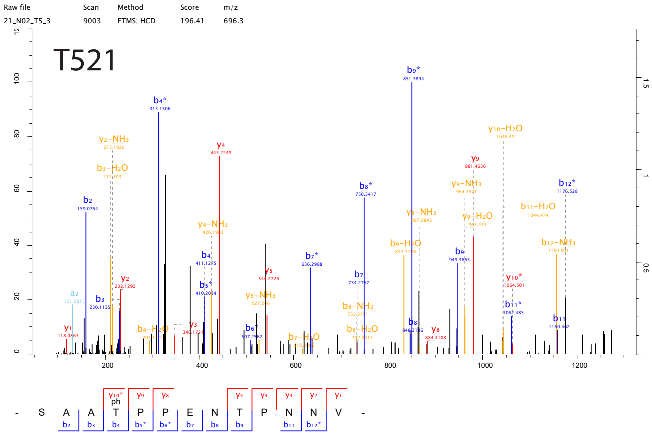

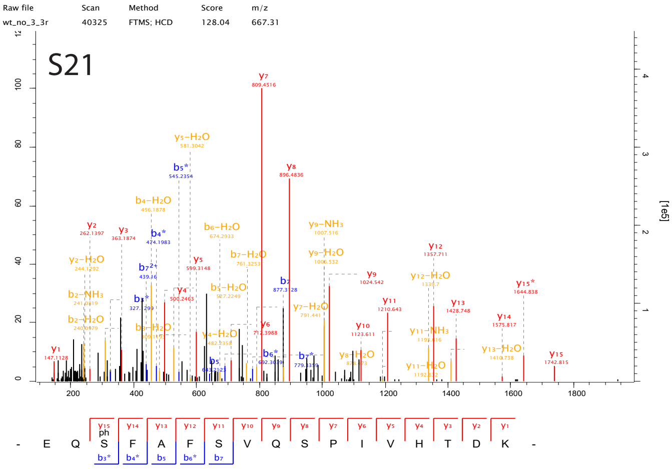

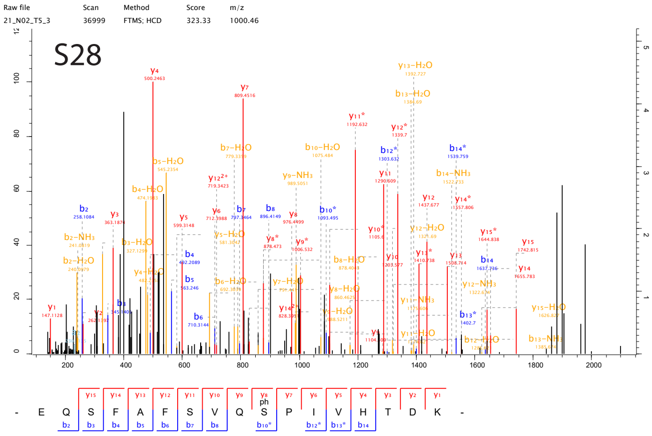

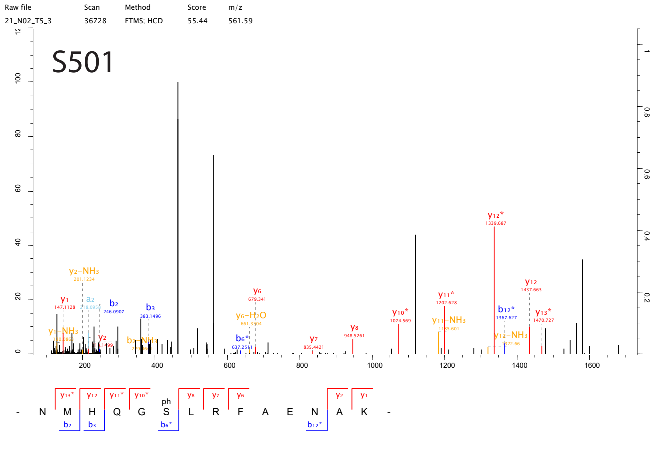

### Supplementary Figure S3

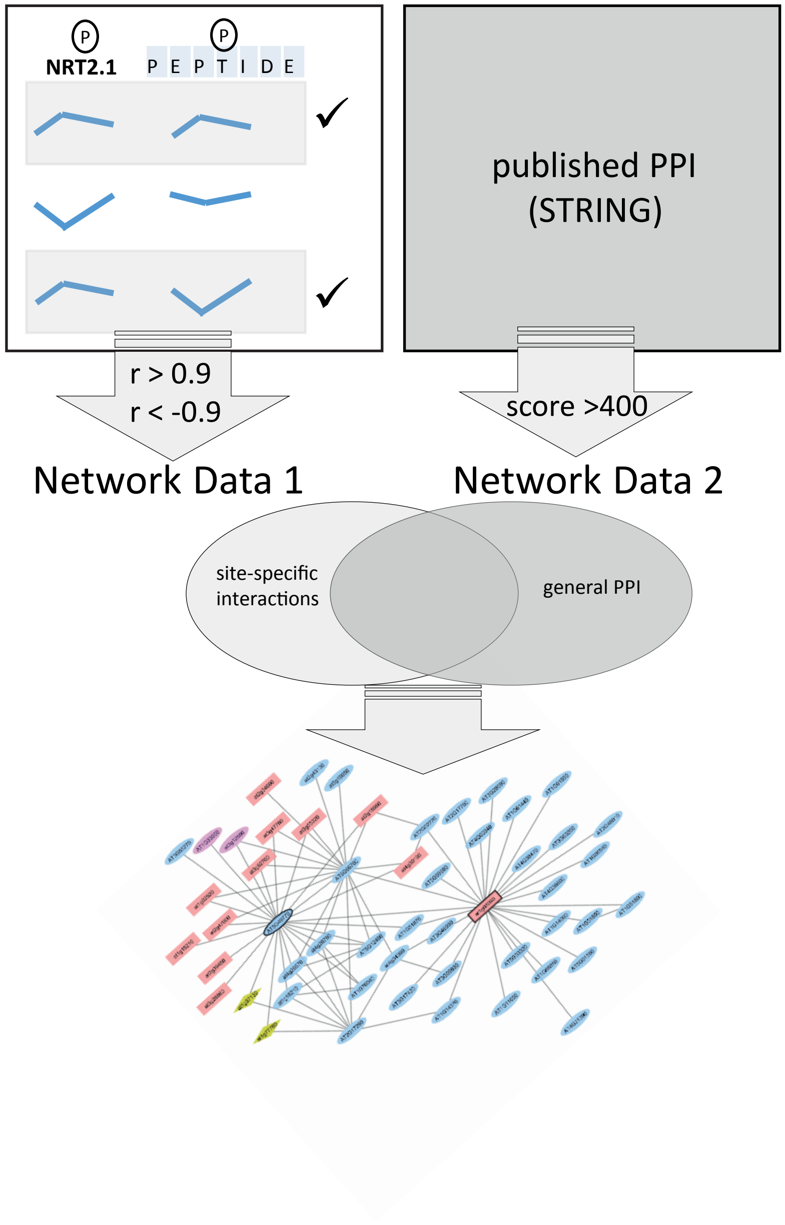

### Supplementary Figure S4

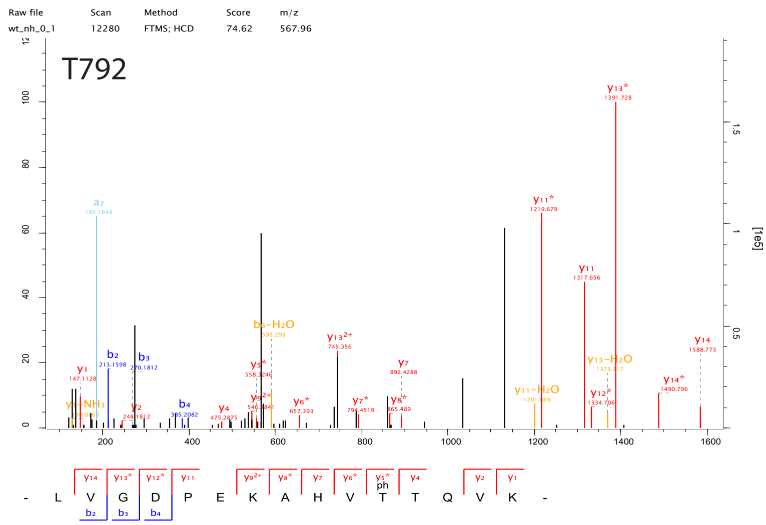

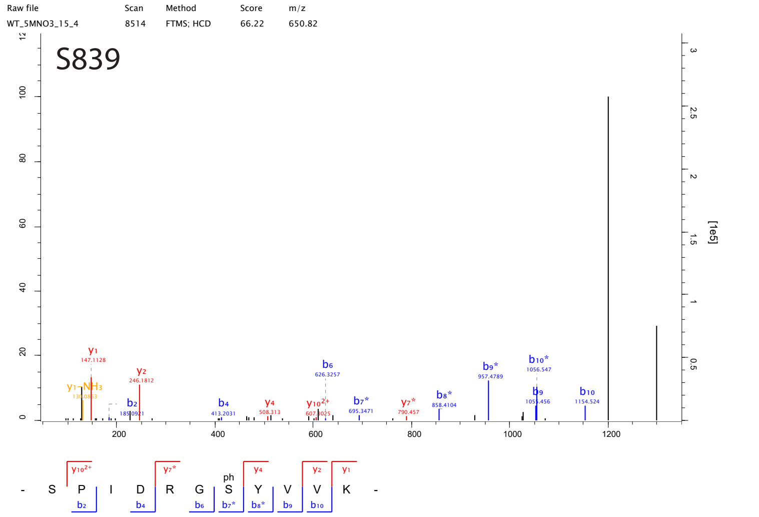

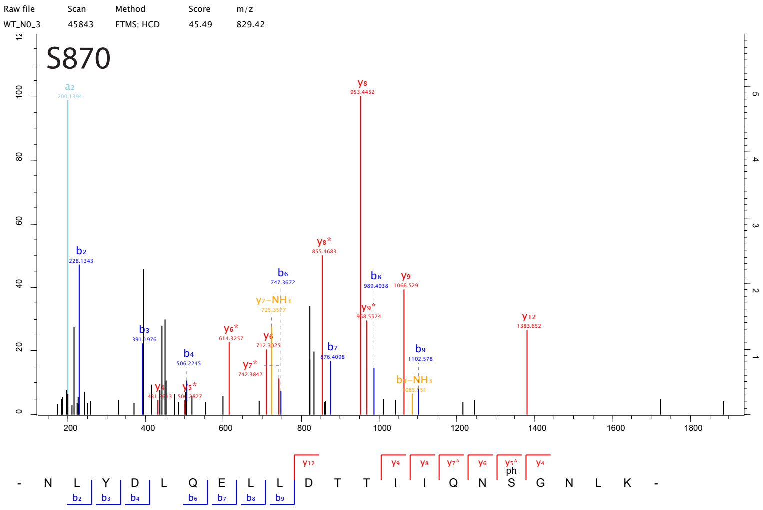

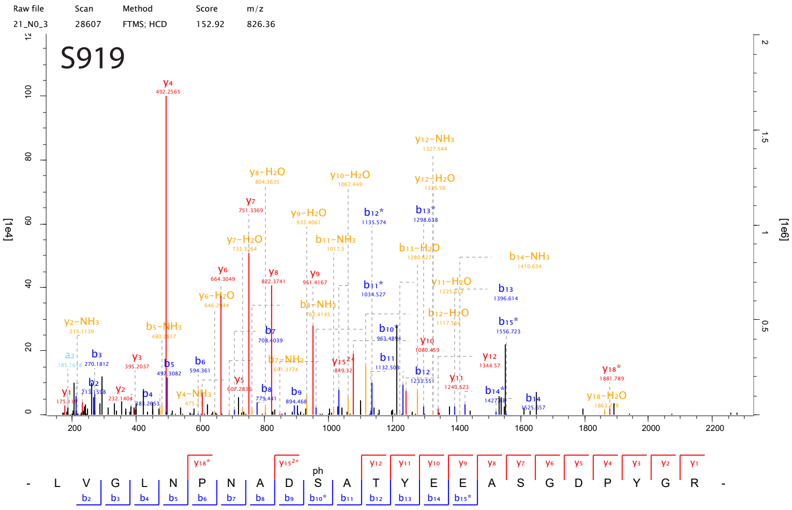

### Supplementary Figure S5

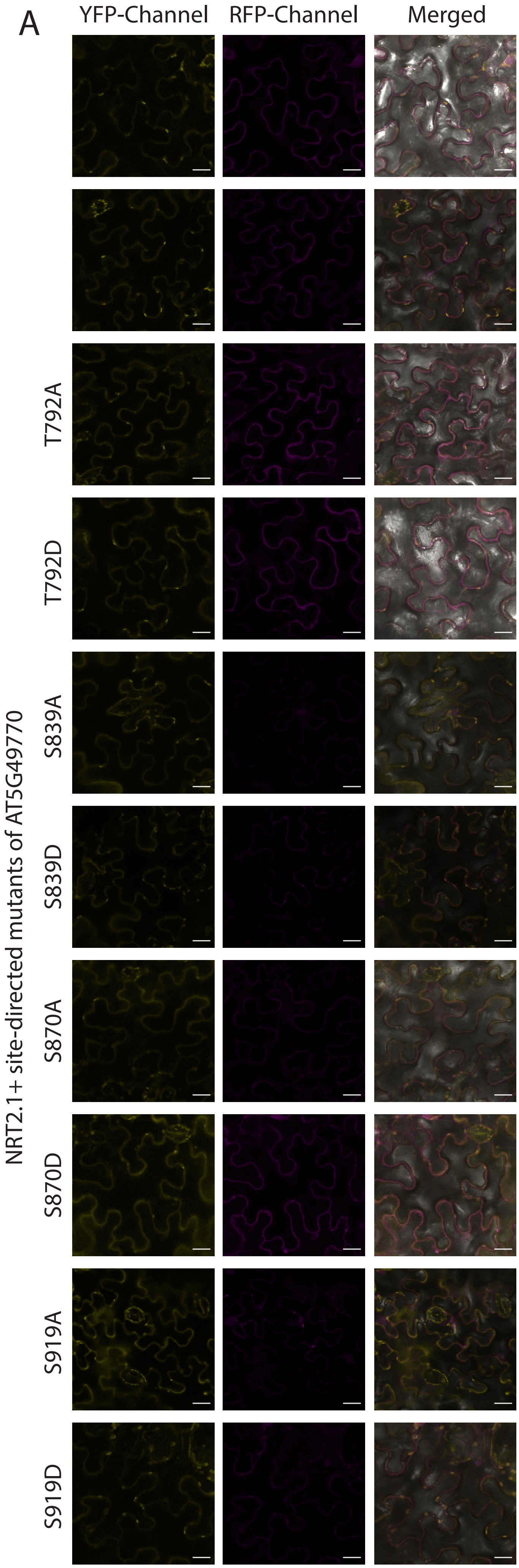

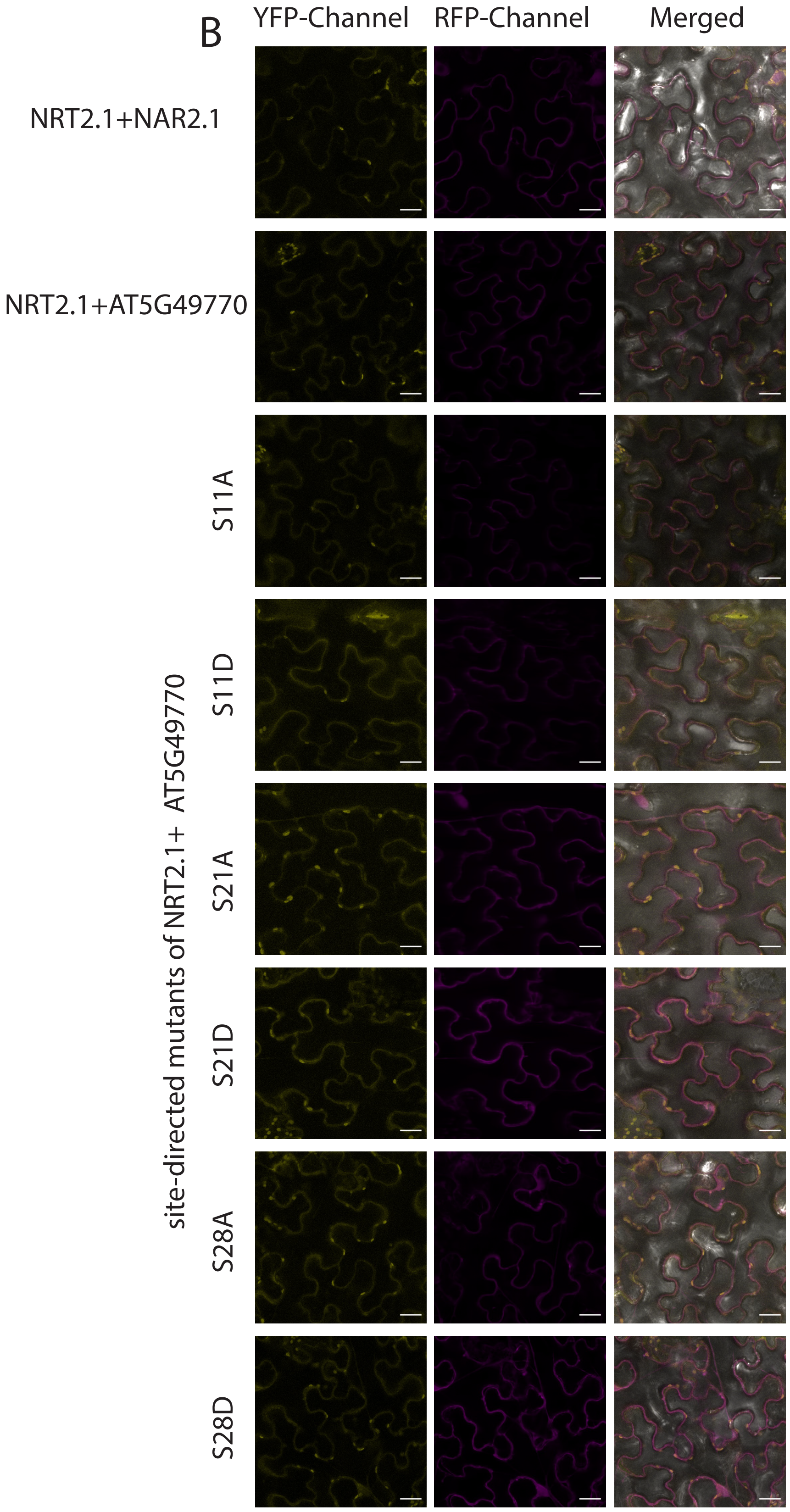

### Supplementary Figure S6

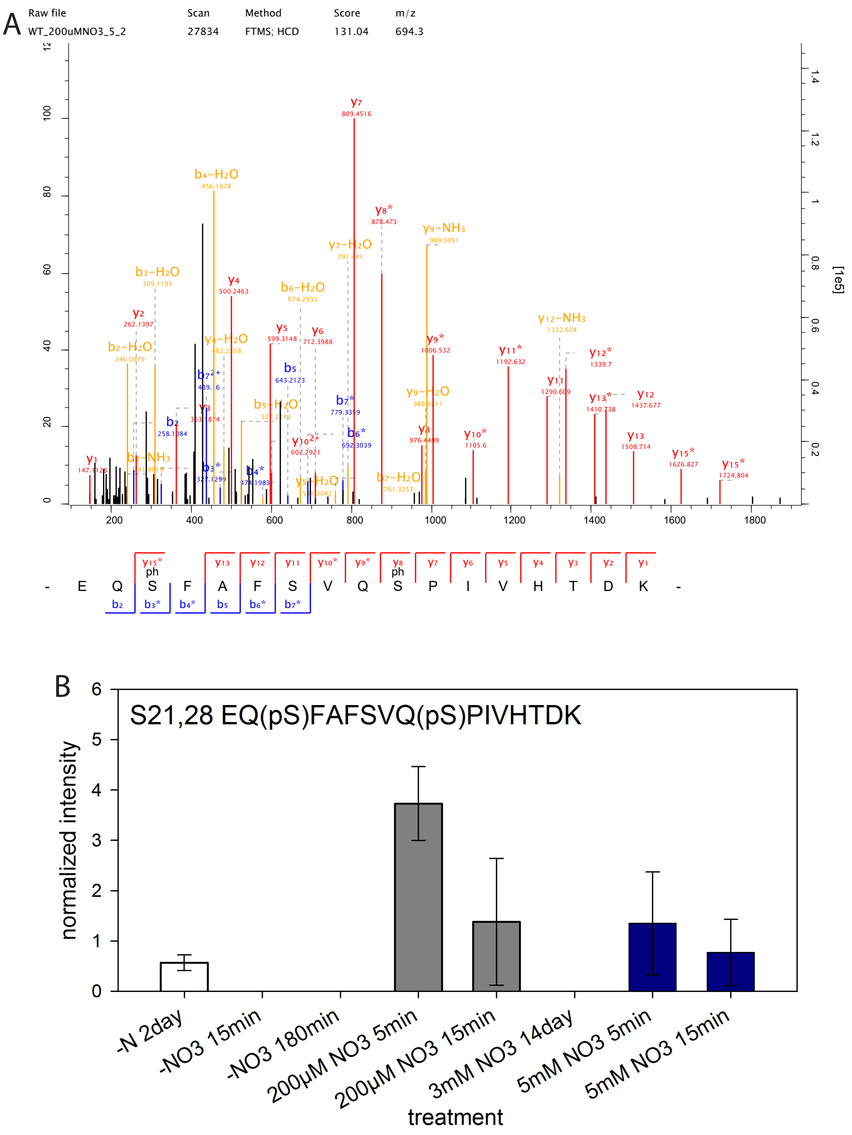

### Supplementary Figure S7

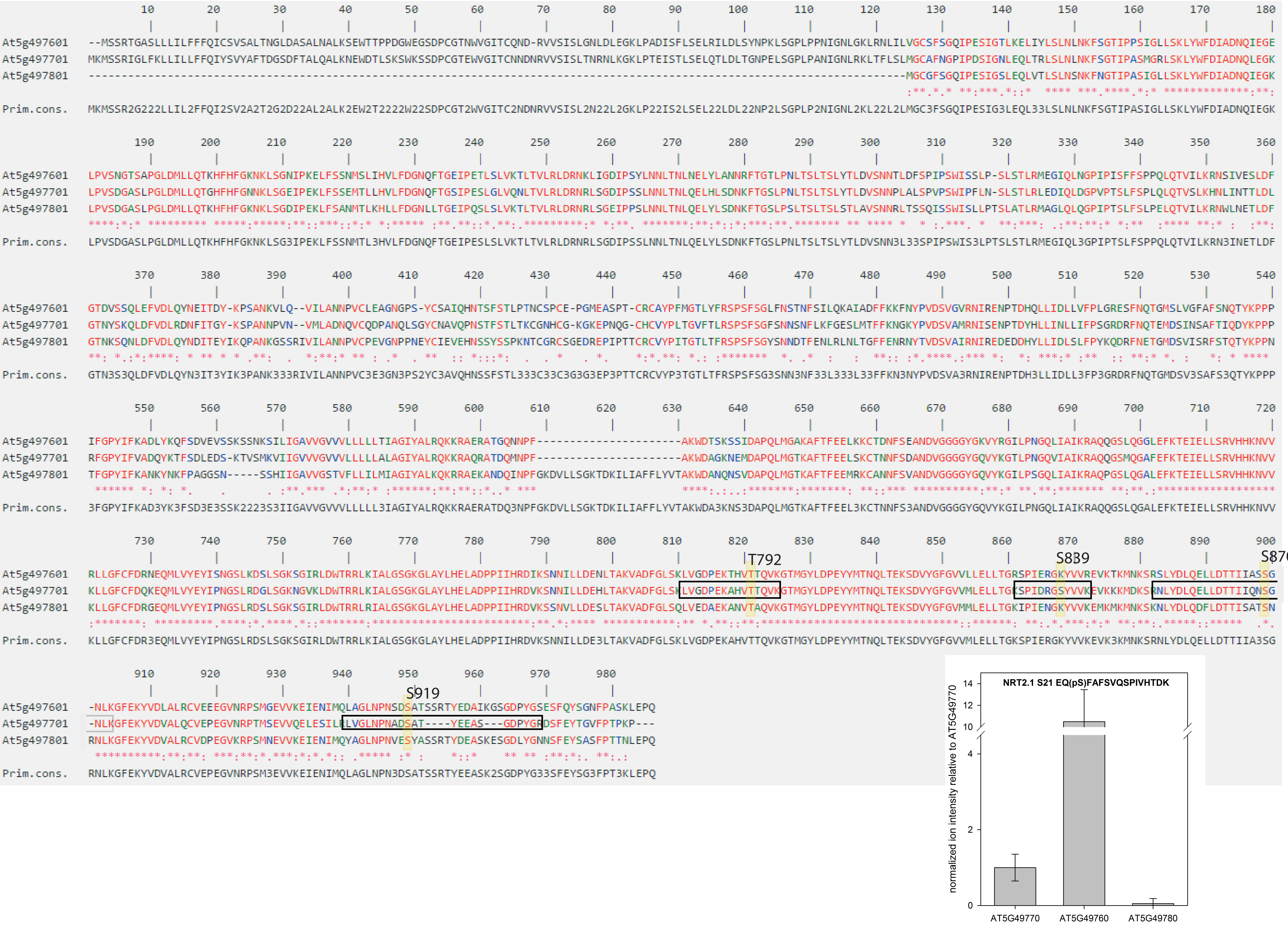
