## Supplementary Material Description for "A phospho-switch in the N-terminus of NRT2.1 affects nitrate uptake by controlling the interaction of NRT2.1 with NAR2.1"

**Figure S1**: Experimental data sets used in this study. (**A**) Nitrate deprivation data set sampled after 3 weeks growth on 3 mM $KNO_3$ and after 15 minutes and 3 hours of nitrate deprivation (Menz et al., 2016). (**B**) Nitrate resupply data set sampled after two days of nitrogen starvation and after resupply of 5 mM or 0.2 mM $KNO_3$ for 5 min and 15 min. Red arrows symbolize the sampling times within each data set. Axes not drawn to scale. (**C**) Processes over-represented as affected by mild nitrate deprivation (dashed edges) and upon resupply with 0.2 mM or 5 mM nitrate (solid edges). Edge color: dashed light gray, 15min; dashed gray, 180min; solid light gray, 5 min; solid dark gray, 15 min. Nodes: 4.1 glycolysis.cytosolic branch; 10.1 cell wall.precursor synthesis; 12.1.1. N-metabolism.nitrate metabolism.NR; 26 misc; 27.1 RNA.splicing; 27.3 RNA.regulation of transcription; 28.1 DNA.synthesis/chromatin structure; 29.2.1 protein.synthesis.ribosomal protein; 29.4.1. protein.posttranslational modification.kinase; 29.5 protein.degradation; 30.2 signalling.receptor kinases; 30.3 signalling.calcium; 30.5. signaling.G-proteins; 31.4 cell.vesicle transport; 34.1.2. transport.p- and v-ATPases.H+-exporting ATPase; 34.16 transport.ABC transporters; 34.18 transport.unspecivied anions; 34.19 transport.Major Intrinsic Proteins; 34.2 transport.sugars; 34.5 transport.ammonium.

**Figure S2**: Best spectra directly exported from MaxQuant version 1.5.3.30 (Cox and Mann, 2008) providing evidence for phosphorylation of NRT2.1 at S11 with peptide (ac) GDSTGEPGSS(ph)MHGVTGR, T521 with peptide VRSAAT(ph)PPENTPNNV, S21 with peptide EQS(ph)FAFSVQSPIVHTDK, S28 EQSFAFSVQs(ph)PIVHTDK, and S501 with peptide NMHQGS(ph)LR. The phosphorylated amino acid is followed by (ph).

**Figure S3**: Workflow of identification of protein kinases which might phosphorylate NRT2.1.

**Figure S4**: Best spectra directly exported from MaxQuant version 1.5.3.30 (Cox and Mann, 2008) providing evidence for phosphorylation of NURK1 (AT5G49770) at T792 with peptide LVGDPEKAHVT(ph)TQVK, S839 with peptide SPIDRGS(ph)YVVK, S870 with peptide NLYDLQELLDTTIIQNS(ph)GNLK and S919 with peptide LVGLNPNADS(ph)ATYEEASGDPYGR. The phosphorylated amino acid is followed by (ph).

**Figure S5**: Representative images of the interaction of ratiometric bimolecular fluorescence assays to detect (**A**) the interaction kinase NURK1 (AT5G49770) and its phosphorylation site mutants with NRT2.1 and (**B**) the interaction of phosphorylation site mutants of NRT2.1 with kinase AT5G49770. Images represent the interaction (YPF-channel) and plasmid expression control (RFP-channel) Scale bar: 50 µm.

**Figure S6**: Doubly phosphorylated peptide of NRT2.1 covering sites S21 and S28. (**A**) Representative fragment spectrum identifying the doubly phosphorylated peptide as directly exported from MaxQuant version 1.5.3.30. (**B**) Ion intensity of doubly phosphorylated peptide at various nitrogen treatment conditions. Averages with standard deviation of three biological replicates are shown.

**Figure S7**: Close homologs of NURK1, AT5G49770. The proteins AT5G49760 and AT5G49780 are closely related to the candidate kinase AT5G49770. The identified phosphopeptides are boxed and the respective phosphorylation sites are shaded in yellow. All identified phosphopeptides of AT5G49770 are proteotypic. All phosphorylation sites except S839 are conserved. S839 is a unique phosphorylation sites within the kinase domain of AT5G49770.

**Supplementary Table 1**: Summary of identified phosphopeptides from the joint analysis of nitrate deprivation experiment (Menz et al., 2016) and nitrate starvation – resupply experiment with 0.2 mM and 5 mM nitrate resupply. In both experimental sets, only root material was analyzed. Quantitative values indicate normalized ion intensity ratios as derived from cRacker (Zauber and Schulze, 2012).

**Supplementary Table 2**: Correlation network of NRT2.1 phosphorylation sites with phosphorylation sites of different kinases under nitrogen deprivation (Menz et al., 2016) and nitrate resupply with 0.2 mM and 5 mM nitrate.
